# Septins mediate cell-cycle-dependent exclusion of Cdc42 GAP Rga4 from growth sites in fission yeast

**DOI:** 10.1101/2024.12.20.629769

**Authors:** Justin McDuffie, Goeun Chang, Maitreyi Das

## Abstract

The polarized growth promoter Cdc42 is inactivated at the growing cell ends during mitosis and reactivated after division. In fission yeast, the Cdc42 inactivator/GAP Rga4 localizes to the cell ends during mitosis and is restricted to the sides in interphase. We show that this cell-cycle-dependent Rga4 localization is septin-dependent. Septins form linear filaments along the sides during interphase and recede to form a medial ring during mitosis. In septin mutants, Rga4 is mobile, homogeneously distributed, and increasingly localizes to the cell ends and cytoplasm, similar to mitotic cells. Accordingly, septin mutants are monopolar with disrupted Cdc42 activation dynamics. This regulation appears to be indirect, via septin-dependent lipid organization on the cortex. We find that the septin cytoskeleton prevents excessive phosphatidylinositol 4,5-bisphosphate (PI(4,5)P_2_) and its upstream kinase Its3 at the plasma membrane. Limiting PI(4,5)P_2_ levels enhances Rga4 puncta at the cortex. Our data describe an unusual form of polarity regulation where the septin cytoskeleton restricts GAP localization via membrane lipid organization.

**Summary:** Cell-cycle-dependent Rga4 localization pattern is controlled by the septin cytoskeleton to promote bipolar growth. In interphase, the septin cytoskeleton at the cell sides restricts Rga4 mobility and localization away from the cell ends to promote proper Cdc42 activation dynamics.

## INTRODUCTION

Establishment and maintenance of cell shape is fundamental to its function. Most cells exhibit a consistent cell shape after every round of cell division, and this ensures their ability to maintain function. Consistent cell shape is not just observed in multicellular organisms that form highly organized tissues and organelles, but even in unicellular organisms such a yeast (Das and Verde, 2013; Galliot and Ghila, 2010; Guillot and Lecuit, 2013; Mitchison and Nurse, 1985). In most eukaryotes, polarized cell growth stops during division and resumes immediately after, to establish consistent cell shape (Lloyd, 2013; McCusker and Kellogg, 2012). It is not well understood how polarized growth resumes promptly after each round of cell division. To understand this process, we use the fission yeast model system *Schizosaccharomyces pombe*. These cells stop polarized growth from the cell ends during division and resume growth once the cells complete cytokinesis (Mitchison and Nurse, 1985). However, we have previously shown that resumption of polarized growth from the cell ends is independent of cytokinesis and is instead cell-cycle-dependent and occurs at a fixed time after mitosis (Rich-Robinson et al., 2021).

Like in many eukaryotic cells, the major regulator of polarized growth in fission yeast is the small GTPase Cdc42 (Das and Verde, 2013; Estravis et al., 2012; Etienne-Manneville, 2004; Johnson, 1999). Cdc42 is activated when Guanine nucleotide exchange factors (GEFs) help it bind GTP (Bos et al., 2007; Sprang, 2001). GTP-bound Cdc42 binds and activates downstream targets to promote polarized growth (Bos et al., 2007). Cdc42 is inactivated when it hydrolyses the GTP to GDP with the help of the GTPase-activating proteins (GAPs) (Bos et al., 2007). Cdc42 in fission yeast has two GEFs, Scd1 and Gef1, and three GAPs, Rga4, Rga6, and Rga3 (Chang et al., 1994; Coll et al., 2003; Das et al., 2007; Gallo Castro and Martin, 2018; Murray and Johnson, 2001; Revilla-Guarinos et al., 2016; Tatebe et al., 2008). Our lab previously showed that during cell division, the GAP Rga4 changes its localization pattern (Rich-Robinson et al., 2021). Rga4 localizes to the cell sides in a punctate manner during interphase. It becomes diffuse along the cell cortex and localizes to the cell tips in dividing cells, where it prevents Cdc42 activation. Once cells complete division, Rga4 resumes its punctate localization away from the cell ends, and growth at the ends resumes. It is not known how Rga4 localization is regulated in a cell-cycle-dependent manner.

Rga4 has a coiled-coil region that is required for its localization to the cell cortex (Das et al., 2007). Cells lacking *rga4* show increased width, since in these mutants, the zone of Cdc42 activation and consequent growth spreads over a larger area at the cell ends (Das et al., 2007; Tatebe et al., 2008). It is not known how Rga4 localization is limited to the cell sides and away from the ends. A recent report shows that Rga4 is removed from the cell ends as a result of membrane flow that occurs due to extensile growth at these ends (Gerganova et al., 2021; Rutkowski et al., 2024). During mitosis, Rga4 is diffuse along the membrane and appears at the cell ends (Rich-Robinson et al., 2021). It is not known what prevents the diffuse Rga4 localization at the cell surface during interphase, restricting it to the sides.

In several eukaryotic cells, the septin cytoskeleton acts as a diffusion barrier along the membrane (Bridges and Gladfelter, 2015; Caudron and Barral, 2009; Kinoshita, 2003). *S. pombe* expresses seven total septin proteins (Spn1-7p), however, only four are involved in the regular cell cycle (Spn1-4p) (An et al., 2004; Berlin et al., 2003; Onishi et al., 2010). The septin cytoskeleton is a hetero-octamer consisting of 2 sets of 4 peptides Spn1-4 (An et al., 2004; Berlin et al., 2003). These peptides form hetero-octamers that organize in a palindromic manner to form larger filaments (Kinoshita, 2003; Spiliotis and McMurray, 2020; Woods and Gladfelter, 2021). The septin cytoskeleton plays a role in cytokinesis, where it forms a ring at the division site. The septin ring acts as the landmark where proteins required for cell separation during cytokinesis are delivered (Berlin et al., 2003; Maddox et al., 2007; Martin-Cuadrado et al., 2005; Singh et al., 2024; Spiliotis and McMurray, 2020). It is not clear if the septin cytoskeleton plays a role during interphase. Cells lacking the septin cytoskeleton show cell separation defects but remain polarized (An et al., 2004; Berlin et al., 2003; Martin-Cuadrado et al., 2005).

Here, we show that the septin cytoskeleton prevents Rga4 diffusion along the cell cortex and restricts it to the cell sides. In cells lacking septin filaments, Rga4 at the cell sides is highly mobile, appears diffuse along the cortex, and increasingly localizes to the cell ends, similar to that observed in mitotic cells. Correspondingly, in cells lacking septins, the Cdc42 activation dynamics are disrupted, resulting in monopolar cells. During interphase, the septin cytoskeleton forms small filaments at the cortex that align along the long axis of the cell. The septin levels along the cortex decrease as the cells enter mitosis, and eventually, a septin ring forms at the end of anaphase B. Around this time, Rga4 localization appears diffuse at the cortex, and active Cdc42 is lost from the cell ends. We find that the mechanism for septin-dependent Rga4 localization is not via direct protein-protein interaction as determined by acceptor photobleach FRET and the GFP-GBP trap method. Instead, we find that upon loss of septin filaments, the lipid organization on the plasma membrane is disrupted with enhanced PI(4,5)P_2_ levels and decreased PI(4)P levels at the cortex. Moreover, in these mutants, the localization of the PI(4)P kinase Its3 is also disrupted. Correspondingly, in mutants lacking Its3 kinase function, Rga4 shows enhanced cortical localization. These findings together suggest a role for the septin cytoskeleton in maintaining proper lipid organization at the cortex, thereby restricting Rga4 localization to the cell sides. These findings show that the septin cytoskeleton maintains cell polarity in a cell-cycle-dependent manner.

## RESULTS

### The septin cytoskeleton restricts Rga4 to the cell sides during interphase

To investigate if the septin cytoskeleton restricts Rga4 localization, we analyzed Rga4-GFP in *spn1^+^*and *spn1*Δ cells. In the *spn1^+^* cells, Rga4-GFP along the cell sides appears punctate in interphase cells and diffuse in dividing cells. In contrast, *spn1*Δ cells show a more diffuse Rga4-GFP localization pattern, regardless of the cell cycle stage (Fig.1A). We quantified the level of Rga4 heterogeneity along the cell sides by measuring the coefficient of variance of Rga4-GFP intensity. As reported earlier, we find that the coefficient of variance of Rga4-GFP is highest in the cells in the G2 phase of the cell cycle and the lowest in the mitotic phase in *spn1^+^*cells (Fig. 1B) (Rich-Robinson et al., 2021). This indicates that Rga4-GFP is more homogeneous along the cell sides in mitotic cells and forms puncta during G2 or interphase. In *spn1*Δ cells, the coefficient of variance in G2 cells is low and is similar to that in mitotic cells (Fig. 1C).

**Figure 1.**
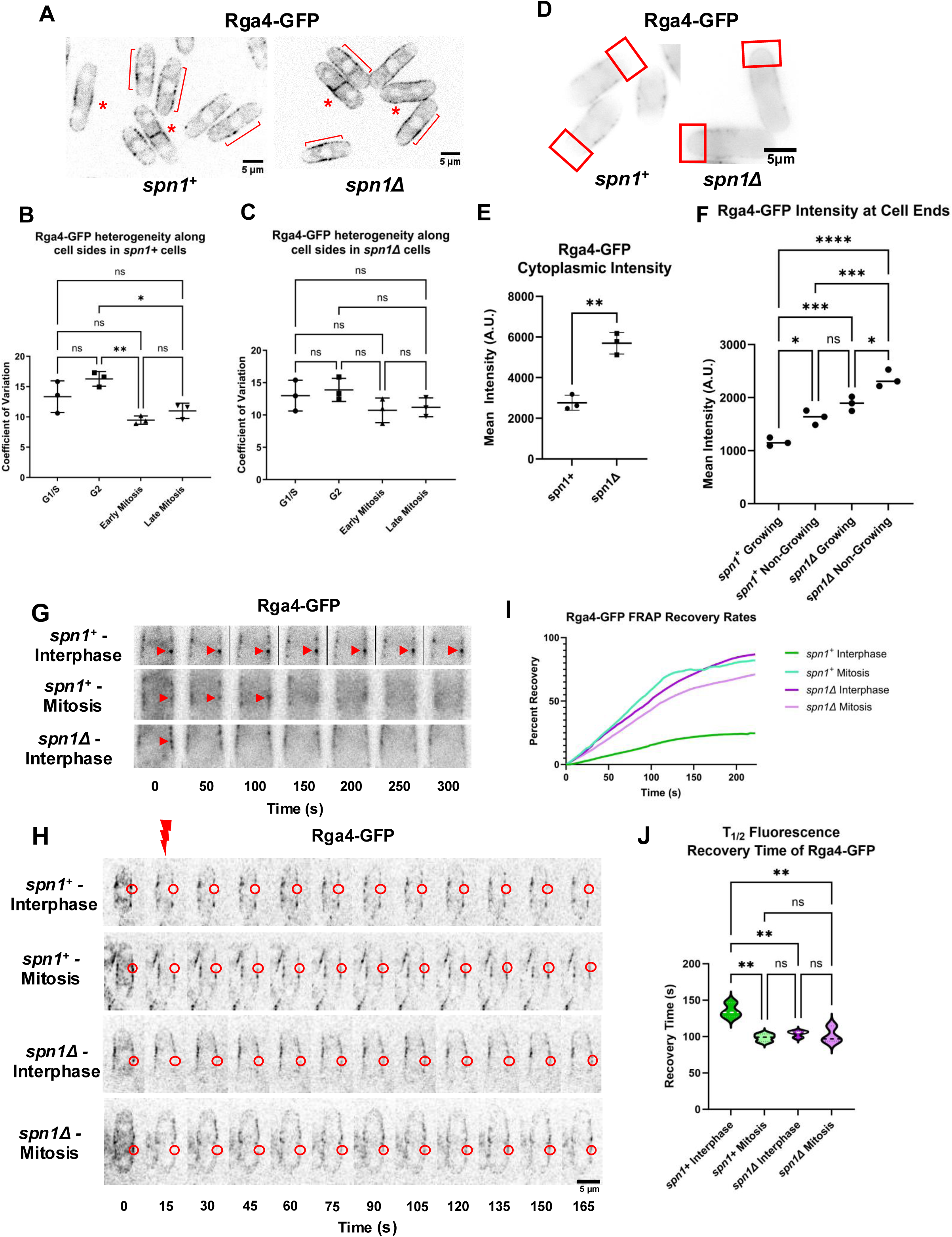
The septin cytoskeleton restricts Rga4 localization and dynamics at the cell sides during interphase. **A.** Middle plane images of Rga4-GFP in *spn1^+^* and *spn1*Δ cells. Red brackets highlight Rga4 distribution along cell sides in interphase cells. Red asterisks highlight dividing cells. **B-C**. Quantification of the coefficient of variation of Rga4 distribution through the cell-cycle in *spn1^+^* (B) and *spn1*Δ (C) cells. [N=3, *n*=40-50 cells each, \**P*≤0.05, *ns*=not significant,]. **D.** Rga4-GFP increasingly localizes to the cell ends and cytoplasm in *spn1*Δ cells during interphase. Images consist of superimposed middle-plane images taken at 10-second intervals over a 5-minute time-lapse. Red boxes highlight increased localization of Rga4 to the cell ends. **E.** Quantification of Rga4-GFP intensity within the cytoplasm of interphase cells in *spn1^+^* and *spn1*Δ conditions measured by averaging three unique regions within the cytoplasm of each cell [ \*\**P*≤0.01, Student’s T test]. **F.** Quantification of Rga4-GFP localization to growing and non-growing cell ends of interphase cells in *spn1^+^* and *spn1*Δ conditions. [N=3, *n*=24-28 cells each, \*\*\*\**P*≤0.0001, \*\*\**P*≤0.001, \*\**P*≤0.01, \**P*≤0.05, *ns*=not significant, one-way ANOVA, scale bar=5μm]. **G.** Montage of Rga4-GFP in *spn1^+^* cells during interphase, during mitosis, and in *spn1*Δ cells during interphase, taken over a 5-minute time-lapse. Red arrowhead marks an Rga4-GFP puncta at the start of the time-lapse. **H.** Montage of Rga4-GFP recovery after photobleaching during interphase and mitosis in *spn1^+^* and *spn1*Δ cells taken in 15-second intervals. Red circles indicate the photobleached region of interest (ROI). Red lightning bolt indicates the time point of photobleaching. **I.** FRAP recovery rates of Rga4-GFP signal intensity within the ROI over time in *spn1^+^* and *spn1*Δ cells during interphase and mitosis. **J.** T_1/2_ fluorescence recovery times of listed conditions. [N=3, *n*=10 cells each, \*\**P*≤0.01, \**P*≤0.05, *ns*=not significant, one-way ANOVA, scale bar=5μm].

In mitotic cells, Rga4 appears more cytoplasmic and is also increasingly observed at the cell ends. We asked if septins restrict cytoplasmic Rga4 and prevent its localization to the cell ends. Rga4-GFP along the cell ends appears faint, even in Z-projected images, due to the curvature of the cell and weak expression/signal. To overcome this, we took rapid time-lapse movies with 5-second intervals over 5 mins of Rga4-GFP expressing *spn1+* and *spn1*Δ cells. We measured Rga4-GFP intensity in the cytoplasm, and the cell ends of time-projected images of these movies (Fig.1D). We find that compared to *spn1^+^*cells, cytoplasmic Rga4-GFP levels increase in *spn1*Δ cells (Fig.1E). Similarly, *spn1*Δ cells show enhanced Rga4-GFP levels at the cell ends compared to *spn1^+^*cells (Fig.1F). In control cells, Rga4-GFP shows elevated levels at the non-growing cell ends compared to growing ends (Gerganova et al., 2021). We also see higher levels of Rga4-GFP at the non-growing cell ends of *spn1*Δ cells, but these levels are much higher than those in *spn1+* cells. Moreover, the growing ends in *spn1*Δ cells show higher Rga4-GFP levels compared to growing ends of *spn1^+^* cells. This indicates that in the absence of septins, Rga4-GFP levels at the cell ends are elevated.

### Septins inhibit Rga4-GFP mobility during Interphase

Next, we asked whether the increased homogeneity of Rga4-GFP along the plasma membrane is the result of increased Rga4-GFP mobility or diffusion along the membrane. The above-mentioned 5-minute time-lapse movies of Rga4-GFP localization were analyzed, and we found that, in *spn1^+^* cells in interphase, Rga4-GFP formed a number of bright, stationary puncta along the plasma membrane that remained immobile throughout the course of time-lapse imaging (Fig. 1G). In contrast, *spn1^+^* cells in mitosis, and *spn1*Δ cells at all stages of the cell-cycle showed limited Rga4-GFP puncta formation, and lacked immobile Rga4-GFP puncta. In these cells, Rga4-GFP appeared to diffuse more rapidly along the plasma membrane. To quantify this observed change in Rga4 mobility, fluorescence recovery after photobleaching (FRAP) microscopy was performed on 10x10-pixel ROIs of Rga4-GFP along the cell sides of *spn1^+^* and *spn1*Δ cells in interphase and mitosis (Fig. 1H). Cell-cycle progression was marked by the fluorescently tagged spindle pole body marker Sad1-mCherry and the fluorescently tagged Myo2 light chain actomyosin ring marker Rlc1-tdTomato. Fluorescence recovery curves and calculated recovery half-lives of Rga4-GFP in the photobleached ROIs showed that Rga4-GFP was much less mobile along the plasma membrane in interphase *spn1^+^* cells (Fig. 1I, J). In contrast, Rga4-GFP is more mobile along the plasma membrane in *spn1^+^* cells during mitosis. Similar to mitotic *spn1^+^* cells, Rga4-GFP is more mobile along the plasma membrane in *spn1*Δ cells, regardless of cell-cycle stage. This indicates that in the presence of septin filaments, the diffusion of Rga4-GFP is significantly restricted, leading to decreased protein mobility. This decreased mobility along the plasma membrane may also explain the decreased localization of Rga4-GFP to the cytoplasm and the cell ends in the presence of septins.

A previous report has indicated that the GAP Rga6 interacts with the septin filaments and is required for proper septin localization at the cortex in interphase cells (Zheng et al., 2022). Thus, we asked if the septin filaments also regulate Rga6 distribution along the cortex. Similar to Rga4-GFP, Rga6-GFP appears punctate along the cell sides (Supplementary Fig. S1A). However, in *spn1*Δ mutants, Rga6-GFP localization remained the same, and the coefficient of variance along the cortex was not significantly distinct between *spn1^+^*and *spn1*Δ cells, and throughout the cell cycle stages (Supplementary Fig. S1B). This indicates that the septin cytoskeleton specifically disrupts Rga4 distribution along the cortex.

### Septins promote bipolar growth, separate from their role in facilitating proper cytokinesis and cell separation

Next, we asked if, in addition to the changes in Rga4 localization, the septin mutants displayed polarity defects. We observe that the growth patterns of *spn1*Δ mutants are altered compared to their *spn1^+^* counterparts. Notably, cells lacking the core septin *spn1* showed a number of septation defects as a result of failure to successfully separate after division (Fig. 2A, C), as has been reported before (An et al., 2004; Perez et al., 2015; Tasto et al., 2003; Wang et al., 2015; Wu et al., 2010; Zheng et al., 2018; Zheng et al., 2024). These septin mutants also show a significant change in their polarity, with 54% of the cells showing a monopolar growth pattern, as compared to *spn1^+^* cells (18%). The monopolar pattern in *spn1*Δ has been reported earlier, and this has been attributed to the cytokinesis defect observed in these cells, possibly making the new ends incapable of growth (An et al., 2004; Bohnert and Gould, 2012) (Fig. 2A, B). Similarly, we find that in the *spn4*Δ mutants, the cells show cytokinesis defects, as indicated by the high septation index and increased monopolarity (Fig. 2D-F). Both Spn1 and Spn4 are the core septin proteins and are necessary to form the septin oligomer (An et al., 2004). Previous reports have shown that Spn2 and Spn3 are not necessary to form the septin oligomer and their loss do not lead to cytokinetic defects (An et al., 2004). However, we find that even in *spn2*Δ and *spn3*Δ mutants, while the septation index is not elevated, they continue to show increased monopolarity similar to that observed in *spn1*Δ and *spn4*Δ mutants (Fig D-F). This suggests that the polarity defect observed in the septin mutants is independent of cytokinetic defects. Taken together, these data show that the absence of any of these four septin monomers results in polarity defects, even when cytokinetic and septation defects are not present.

**Figure 2.**
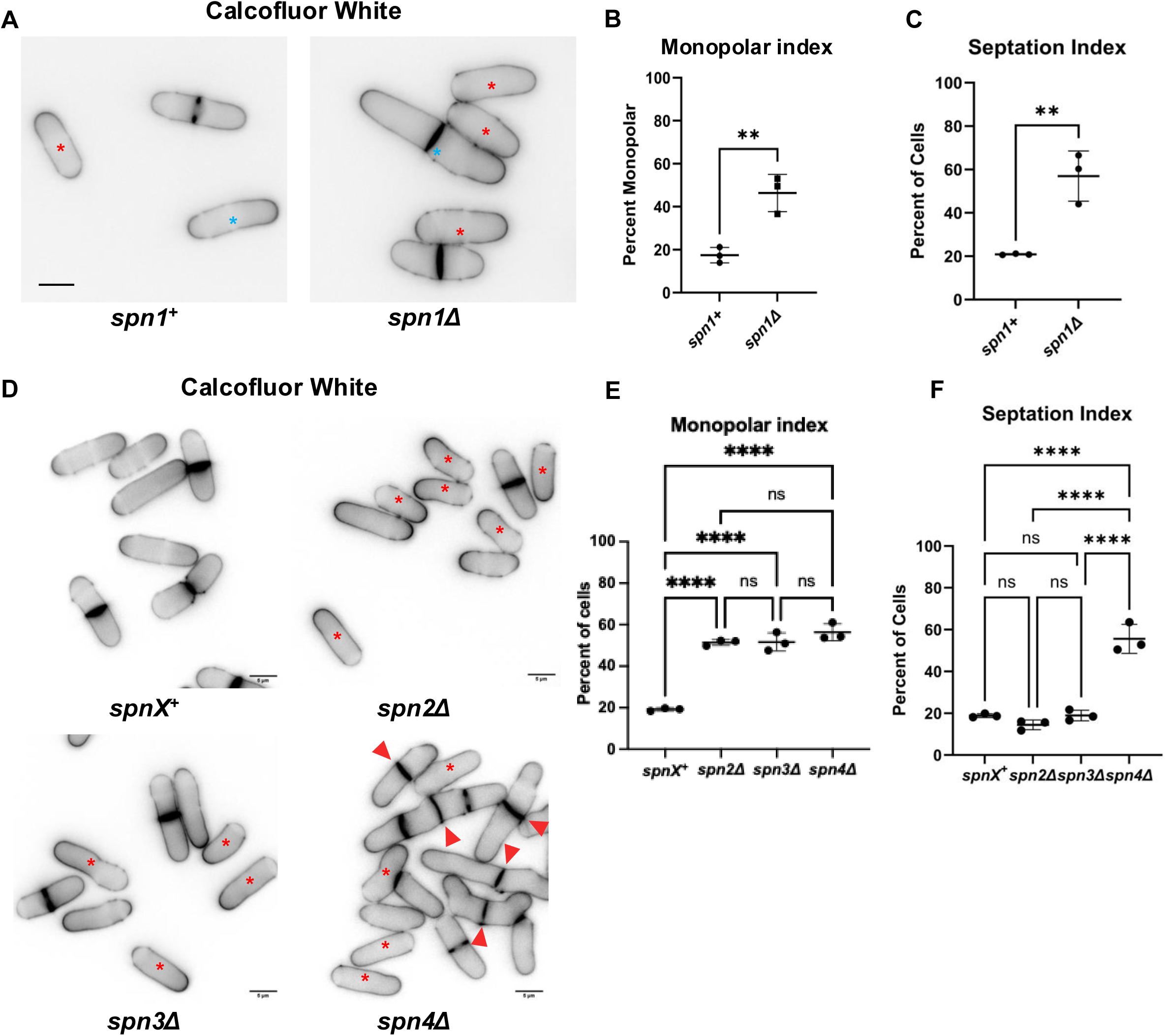
The septin cytoskeleton promotes bipolar growth. **A.** Representative images of calcofluor white-stained *spn1^+^*and *spn1*Δ cells. Blue asterisks highlight bipolar (*spn1^+^*) growth and red asterisks highlight monopolar (*spn1*Δ) growth. **B.** Quantification of monopolarity index in *spn1^+^*and *spn1*Δ cells. [N=3, *n*=65-100, \**P*≤0.05, \*\**P*≤0.01, student’s t-test, scale bars=5μm]. **C.** Quantification of the septation index of *spn1^+^*and *spn1*Δ cells. [N=3, *n*>200, \**P*≤0.01, student’s t-test]. **D.** Representative images of calcofluor white-stained *spnX^+^* and *spn2*Δ*, spn3*Δ, and *spn4*Δ cells, respectively. Red asterisks highlight monopolar cells. Red arrowheads highlight septated cells that failed to separate. **E.** Quantification of monopolarity index in *spnX^+^*and *spn2*Δ*, spn3*Δ, and *spn4*Δ cells, respectively. **F.** Quantification of the septation index of *spnX^+^* and *spn2*Δ*, spn3*Δ, and *spn4*Δ cells, respectively. [N=3, *n=*>110 cells each, \*\*\*\**P*≤0.0001, ns=not significant, one-way ANOVA, multiple comparison test, scale bars=5μm].

### Septins promote proper Cdc42 oscillatory dynamics

Our data show that in the absence of *spn1*, Rga4 localizes to the cell ends, and these cells are mostly monopolar. Given that Rga4 is a GAP for Cdc42, we asked if the septin cytoskeleton impacts Cdc42 activation levels at the cell ends. We find that neither the loss of *spn1*, or *rga4*, or both, result in significant changes to the average levels of Cdc42 activity at the growing ends of cells, as measured using the active Cdc42 bio-marker, CRIB-3xGFP (Fig. 3A, B).

**Figure 3.**
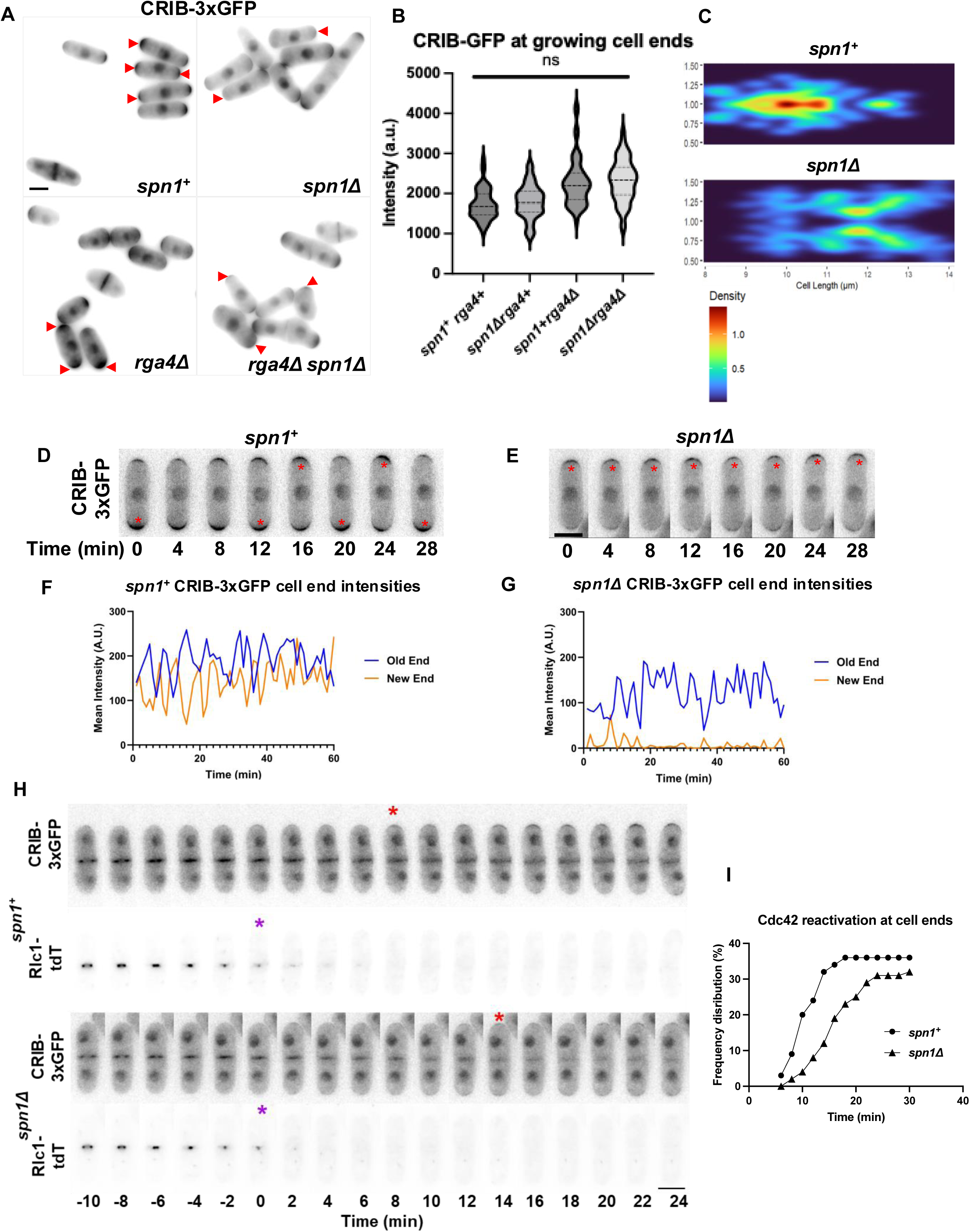
The septin cytoskeleton promotes bipolar Cdc42 activation. **A.** Representative SUM Z-Projection images of *rga4^+^ spn1^+^*, *rga4^+^ spn1*Δ*, rga4*Δ *spn1^+,^* and *rga4*Δ *spn1*Δ cells expressing CRIB-3xGFP, respectively. Red arrowheads highlight Cdc42 activity at the ends of growing cells. **B.** Quantification of the total levels of CRIB-3xGFP intensity via measuring integrated density at growing cell ends of the aforementioned cells in A [N=3, *n*>50, ns=not significant, one-way ANOVA, multiple comparison test, scale bar=5μm]. **C.** Heat map of the fraction of CRIB-GFP intensity at each cell end over the total intensity with increasing cell size in *spn1^+^* and *spn1*Δ cells. **D, E.** Representative images of the oscillatory dynamics of Cdc42 activity via the active Cdc42 biomarker, CRIB-3xGFP, over time in *spn1^+^*and *spn1*Δ cells. Images taken at 1-minute intervals over a 60-minute time-lapse. **F, G.** Quantification of CRIB-3xGFP intensity at the old and new cell ends of a representative *spn1^+^* and *spn1*Δ cell over 60-minute time-lapse**. H.** A representative time-lapse of CRIB-3xGFP and Rlc1tdTomato in *spn1^+^*and *spn1*Δ cells. Red asterisk marks the onset of stable CRIB-3xGFP localization at the cell ends after the completion of ring constriction marked with a purple asterisk. **I.** Frequency plot of the timing of the onset of stable Cdc42 activity at the cell ends as indicated by CRIB-3xGFP after the completion of ring constriction in *spn1^+^* and *spn1*Δ cells (*n*>30). [scale bars=5μm].

As reported previously, an analysis of CRIB-3xGFP fraction at the ends of *spn1^+^* cells appears symmetric between the two ends as the cells increase in size beyond 9μm (Fig. 3C) (Das et al., 2012). However, in *spn1*Δ cells, the ends show increased asymmetry of the CRIB-3xGFP fraction between the two cell ends, and symmetry is partially achieved only at a much higher cell length, beyond 11μm, just before the cells start to divide (Fig. 3C). These data suggest that in the absence of the septin cytoskeleton, Cdc42 activation dynamics between the two ends is disrupted and accumulates only at one end, resulting in monopolarity. Next, we analyzed Cdc42 oscillatory dynamics at the cell ends using the CRIB-3xGFP probe in *spn1^+^* and *spn1*Δ cells. As reported earlier, CRIB-3xGFP shows anticorrelated oscillations between the two cell ends in control cells (Das et al., 2012). However, in *spn1*Δ cells, CRIB-3xGFP activity fluctuated only at the growing end, while the non-growing end barely showed any signal (Fig. 3D-G). While some activation of Cdc42 was transiently observed at the non-growing end, it was swiftly deactivated before establishing a competitive draw for the Cdc42 activators at the growing cell end.

We have previously shown that precocious Cdc42 activity and consequent polarized growth at the cell ends require both the loss of *rga4* and the activation of the MOR/Orb6 pathway (Gupta et al., 2014; Ray et al., 2010; Rich-Robinson et al., 2021). Cdc42 is reactivated at the cell ends as cytokinesis completes to enable old end growth (Rich-Robinson et al., 2021). We asked if Cdc42 reactivation at the ends is delayed in *spn1*Δ mutants. Indeed, we find that in *spn1^+^*cells, Cdc42 is reactivated at the cell ends about 10 minutes after the completion of actomyosin ring constriction as indicated by the probe CRIB-3xGFP and ring marker Rlc1-tdTomato (Fig. 3H, I). In *spn1*Δ mutants, Cdc42 reactivation at the ends occurs later, about 15 minutes after the completion of actomyosin ring constriction (Fig. 3H, I). This suggests that Cdc42 activity at the cell ends is disrupted in the absence of the septin cytoskeleton. These findings indicate that the septin cytoskeleton regulates cell-cycle-dependent Rga4 localization, along with proper Cdc42 activation dynamics at the cell ends.

### The septin cytoskeleton shows distinct localization patterns during interphase and mitosis

Given that the septin cytoskeleton promotes proper cell polarization regardless of a cytokinetic defect, we next investigated how septins organize within the cell in interphase as the cell grows. Previous reports have shown that in interphase, the septin cytoskeleton forms small filaments along the plasma membrane (An et al., 2004; Gregory et al., 2025; Zheng et al., 2018). However, epitope tagging of individual septin proteins, such as Spn1 with tdTomato, shows some functional deficiencies (Gregory et al., 2025). To determine the proper organization of the septin proteins, we first investigated the localization patterns of the individual septin proteins during interphase. We find that Spn1-mEGFP shows very faint cortical puncta during interphase, as has been reported earlier (Fig.4A) (Gregory et al., 2025). With a brighter fluorophore, tdTomato, Spn1 appears to localize to the plasma membrane as short linear filaments along the long axis of the cell (Fig. 4A). Similar short filaments were also visible with Spn2-GFP, Spn3-GFP, and Spn4-tdTomato (Fig. 4A).

**Figure 4.**
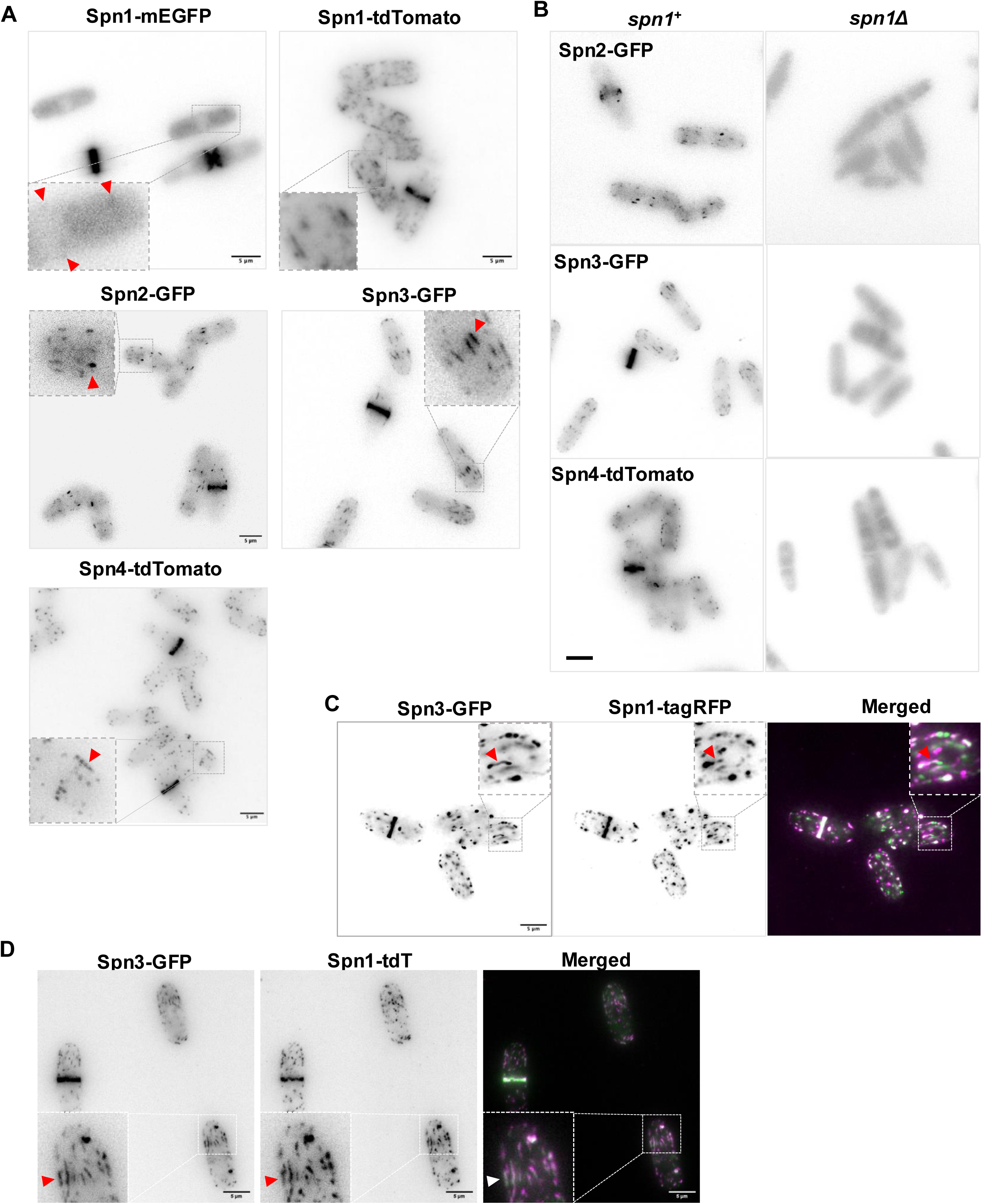
Septins form linear filaments along the cell sides in interphase. **A.** Localization patterns of Spn1-mEGFP, Spn1-tdTomato, Spn2-GFP, Spn3-GFP, and Spn4-tdTomato. All septin proteins localize to the cell sides as linear filaments (red arrowhead) and form a ring at the division site during mitosis. **B.** Spn2-GFP, Spn3-GFP, and Spn4-tdTomato fail to localize to the cell sides in *spn1*Δ mutants. **C.** Spn3-GFP and Spn1-tagRFP colocalize to the short linear filaments along the cell sides in interphase and within a ring during division. **D.** Spn3-GFP and Spn1-tdTomato colocalize in linear filaments along the cell cortex during interphase, and within a ring during division. Red and white arrowheads highlight filaments. Scale bar=5μm.

We next asked if these interphase filaments were dependent on the core septin protein Spn1. We find that in the absence of *spn1*, Spn2-GFP, Spn3-GFP, and Spn4-tdTomato are cytoplasmic and fail to localize to the plasma membrane (Fig.4B). These proteins also fail to form the septin ring in dividing cells, as has been reported before. This indicates that the core septin protein Spn1 is required for septin filament formation in both interphase and mitosis. However, the filaments are distinctly organized during these two phases.

Next, we asked if the septin proteins colocalize to these filaments on the plasma membrane in interphase. Previous reports suggest that Spn1 proteins tolerate tagging with monomeric fluorophores better than dimeric tags (Gregory et al., 2025). Thus, we first investigated Spn1-tagRFP and Spn3-GFP colocalization in interphase. We find that both Spn3-GFP and Spn1-tagRFP colocalize to the linear filaments along the long axis of the cell (Fig. 4D). We next verified this interaction using strains co-expressing Spn1-tdTomato and Spn3-GFP. Spn1-tdTomato and Spn3-GFP also co-localize as short linear filaments along the cell cortex (Fig. 4D).

We also find that septin filament distribution appears biased towards the growing cell ends. To confirm this, we investigated Spn3-GFP localization in *tea1+* and *tea1*Δ mutants. The *tea1*Δ mutants are monopolar and only grow from one end (Mata and Nurse, 1997; Verde et al., 1995). We find that in *tea1*Δ cells, Spn3-GFP filaments appear closer to the growing cell ends (Supplementary Fig. S2A, red arrowheads). In *tea1*Δ cells, Spn3-GFP levels are higher at the growing ends compared to the *tea1+* cells (Supplementary Fig. S2B). This is likely due to the fact that in *tea1+* cells, Spn3-GFP is distributed between the two growing ends of bipolar cells, while in *tea1*Δ cells, all of the Spn3-GFP accumulates at the single growing end.

Since the septin cytoskeleton appears to restrict Rga4 mobility, we asked whether the septin proteins themselves were mobile along the cell cortex. To test for mobility, we performed FRAP analysis of Spn1-tdTomato along the cell sides in both interphase and dividing cells, and analyzed the fluorescence recovery over a 5-minute timespan. Regardless of the cell-cycle stage, we barely see any recovery of the fluorescent signal 5 minutes post-photobleaching of Spn1-tdTomato (Supplementary Fig. S3A & B). This indicates that the septin filaments observed along the cell sides are stable and relatively immobile.

### Septin filaments recede from the cell sides as the cell enters mitosis

Our data show that Rga4 localization at the cortex is mostly punctate and immobile in interphase and diffused and mobile in dividing cells. In addition, we find that the septins show short filaments along the cell cortex in interphase, and this localization is lost once the cell enters mitosis and a septin ring forms at the division site. Based on this, we propose that Rga4 is more dynamic at the cortex when the septins are lost from this region. To verify this hypothesis, we observed Rga4-GFP and Spn1-tdTomato localization along the cortex in cells at different cell-cycle stages (Supplementary Fig.S4A). As expected, we find that Rga4-GFP appears punctate along the cell sides in early and late G2 cells. Spn1-tdTomato also localizes to this region. In early and late mitotic cells, Rga4-GFP appears more diffuse, and Spn1-tdTomato levels along the sides diminish. Given the low expression levels of Rga4, it was not possible to image Rga4-GFP by time-lapse microscopy throughout the different cell-cycle stages along with Spn1-tdTomato. The fully matured Spn1-tdTomato ring appears in post-mitotic cells while Rga4-GFP shows diffuse localization in mitotic cells as the septin localization along the cortex decreases (Supplementary Fig.S4A, early and late mitosis). This suggests that the removal of septins from the cell cortex, and not the formation of the septin ring, is sufficient to enable diffuse Rga4 localization.

Cdc42 is active at the cell ends in interphase and at the division site in dividing cells (Merla and Johnson, 2000; Wei et al., 2016). Our previous data show that Cdc42 activation is lost at the ends during mitosis and is activated at the division site when the cell forms the cytokinetic actomyosin ring. Moreover, delaying Cdc42 activation at the division site did not change the timing of Cdc42 inactivation at the cell ends (Wei et al., 2016). This indicates that Cdc42 inactivation at the ends is independent of its activation at the division site. We analyzed the timing of CRIB-mCherry loss from the cell ends with respect to mitotic progression, as indicated by Sad1-mCherry and changes in septin Spn3-GFP localization (Fig. 5A). We find that CRIB-mCherry is lost from the cell ends about 10-20 minutes after the onset of mitosis. In contrast, the septin ring forms at the end of Anaphase B (Fig. 5B). This suggests that as the cells enter mitosis, septin levels at the cell sides drop, Rga4 localization changes, and this coincides with the loss of active Cdc42 from the cell ends.

**Figure 5.**
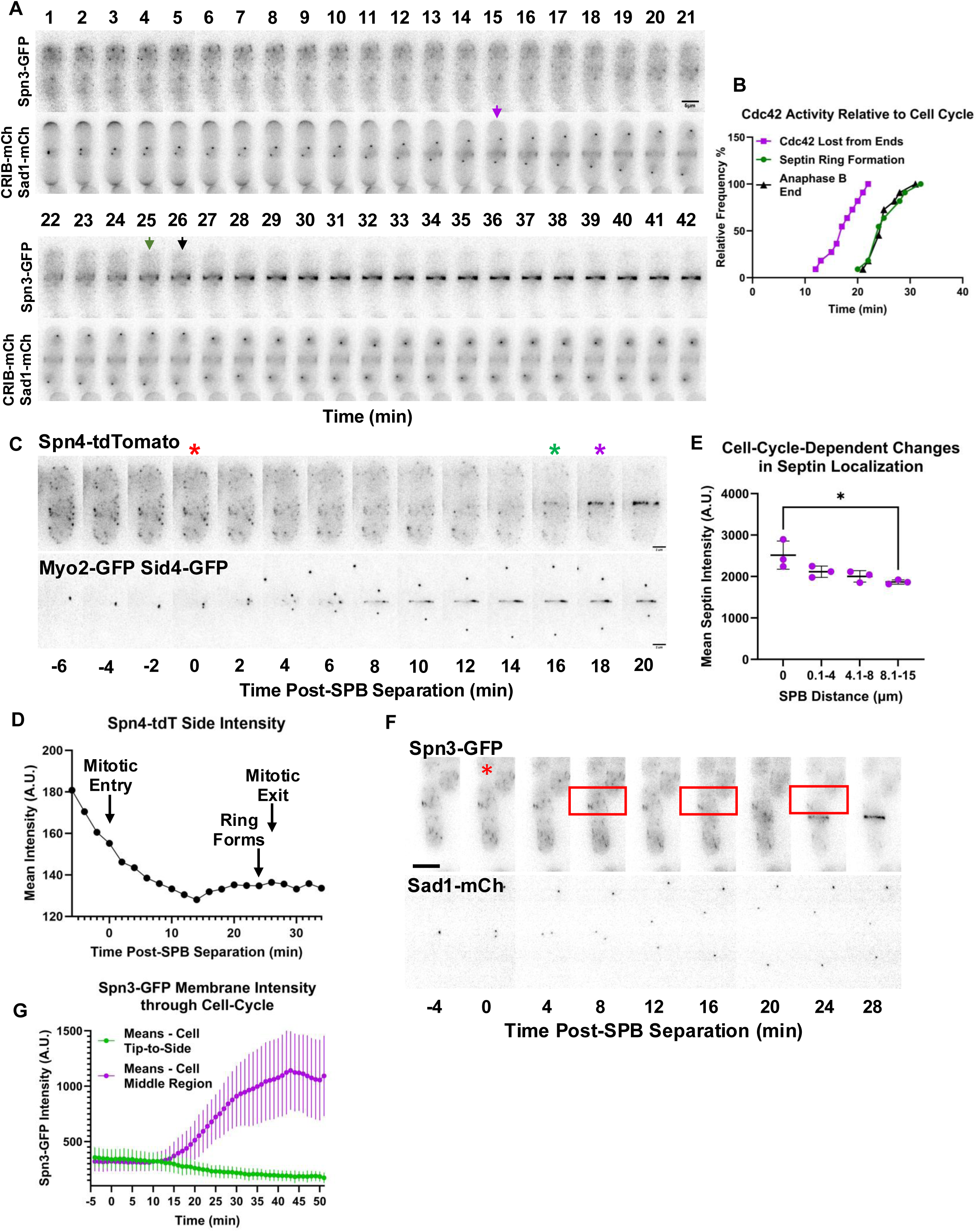
Septins recede from the cell sides during mitosis to eventually form a medial ring as Anaphase B ends. **A.** Representative time-lapse of Spn3-GFP, CRIB-mCherry, and Sad1-mCherry expressing cells starting at the onset of mitosis. Purple arrow highlights the time point when Cdc42 activity is totally lost from the cell ends. Green arrow highlights the maturation of the septin ring. Black arrow highlights the end of Anaphase B. **B.** Frequency plot of the timing of loss of active Cdc42 (CRIB-mCherry) from the cell ends, septin ring formation, and end of Anaphase B with respect to the onset of mitosis. (n=10). **C.** Montage of time-lapse of Spn4-tdTomato and Myo2-GFP Sid4-GFP expressing cells during mitosis. Red asterisk marks the onset of mitosis, the green asterisk marks the onset of septin ring formation, and the purple asterisk marks the end of mitosis. Scale bar=2μm. **D.** Quantification over time of Spn4-tdTomato levels along the cell side of the cell, shown in **C**. **E**. Quantification of Spn4-tdTomato along the cell sides with increasing spindle pole body distances. [N=3, n>20, \**P*≤0.05, one-way ANOVA, multiple comparison test]. **F.** Montage of time-lapse of Spn3-GFP and Rlc1-tdTomato Sad1-mCherry expressing cells during mitosis. Red asterisk marks the onset of mitosis. Red boxes highlight decreasing Spn3-GFP filament intensity along the cell side throughout mitosis. **G.** Quantification of Spn3-GFP intensity along the cell end and sides compared to the cell middle throughout mitosis. [N=3, *n*=10-19, \*\*\*\**P*≤0.0001, student’s t-test]. Scale bar=5μm.

Next, we investigated how septin localization changes throughout the cell cycle. We first imaged cells expressing Spn4-tdTomato along with Myo2-GFP and Sid4-GFP over time-lapse (Fig. 5C). We analyzed Spn4-tdTomato as this is one of the core septin proteins, along with Spn1, required for timely cytokinesis and proper cell separation. While Sid4-GFP labels the spindle pole body and helps visualize the onset of mitosis when the spindle pole body splits into two, Myo2-GFP labels the cytokinetic actomyosin ring (Sparks et al., 1999; Wei et al., 2017). We find that Spn4-tdTomato intensity along the cell cortex drops as the cell enters mitosis, and eventually plateaus (Fig.5D). The Spn4-tdTomato eventually forms a septin ring just prior to mitotic exit, 18 minutes after spindle pole body separation. We quantified Spn4-tdTomato levels along the sides of cells in different mitotic stages, as determined by the distance between spindle pole bodies. We find that Spn4-tdTomato levels along the cell sides are highest in interphase cells with a single spindle pole body (distance, 0μm, Fig. 5E). As the spindle pole body distance increases, Spn4-tdTomato levels along the cell sides drop.

We further looked at Spn3-GFP localization throughout the cell cycle, as its localization is more easily visualized through fluorescence microscopy. Similar to Spn4-tdTomato, Spn3-GFP distribution along the cell sides also begins to change as cells enter mitosis. Septins, which once shared a filamentous localization pattern along the cell sides during interphase, increasingly disappear from these sides upon the onset of mitosis, starting approximately 14 minutes after spindle pole body separation (Fig. 5F, G). At this same time, Spn3-GFP localization to the division plane of the cell increases as the septin ring begins to coalesce and mature (Fig. 5H). This maturation appears to occur approximately 20 minutes after spindle pole body separation, as the spindle poles relax, signaling the end of Anaphase B. Together, these data indicate that septins form short linear filaments at the cortex along the long axis of the cell during interphase. These filaments begin to disappear when the cell enters mitosis, but coalesce into a ring at the division plane only at the end of Anaphase B.

### The septin cytoskeleton and Rga4 show distinct structures at the cortex

Since the Rga4 distribution pattern along the cortex is septin-dependent, we investigated whether the septin cytoskeleton colocalizes with Rga4. We observed Rga4-GFP puncta along the cell sides in cells expressing Spn1-tdTomato. While both these proteins localize to the cell sides, we did not see any significant colocalization pattern between them (Fig.6A, Top Plane, inset). In most cells, Rga4-GFP organizes like a corset at the cell middle as has been reported before (Das et al., 2007; Rich-Robinson et al., 2021; Tatebe et al., 2008), while Spn1-tdTomato appears on either side of Rga4-GFP closer to the growing ends (Supplementary Fig.S4A). In some cells, Rga4-GFP puncta appeared on either side of Spn1-tdTomato filaments (Fig.6A, Middle Plane, inset). To further verify this, we also analyzed cells co-expressing Rga4-GFP and Spn1-tagRFP. We find that Spn1-tagRFP localizes on either side of the Rga4-GFP corset-like organization in interphase cells (Fig. 6B). We asked if the dynamic Rga4-GFP is trapped by septins via physical interaction, thereby preventing its mobility along the cortex.

**Figure 6.**
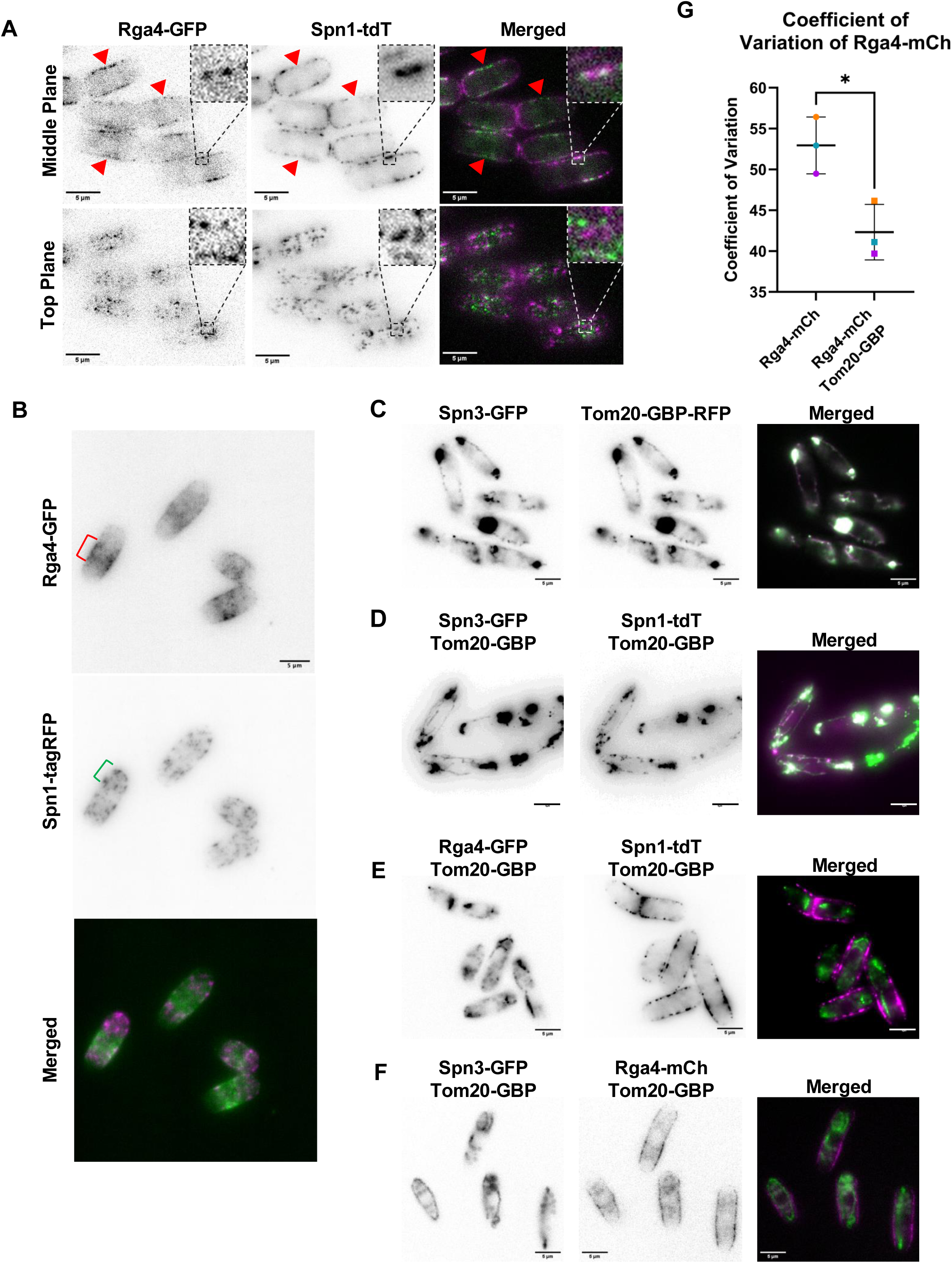
The septin cytoskeleton does not colocalize or interact with Rga4. **A**. Rga4-GFP and Spn1-tdTomato show distinct localization patterns along the sides of interphase cells. Red arrowheads highlight zones with Rga4-GFP but no Spn1-tdTomato along the cell side. The middle plane inset shows partial colocalization of Rga4-GFP with Spn1-tdTomato. The top plane inset shows distinct localization patterns of these proteins. **B.** Rga4-GFP and Spn1-TagRFP show distinct localization patterns in interphase cells. Rga4-GFP localizes as a corset at the cell middle (red bracket) while Spn1-tagRFP appears closer to the cell ends (green bracket). The image is a SUM-projection of a z-stack. **C.** Spn3-GFP mislocalizes to Tom20-GBP-RFP at the mitochondria. **D.** Spn3-GFP and Spn1-tdTomato simultaneously mislocalize to the mitochondria in Tom20-GBP expressing cells. **E.** Rga4-GFP mislocalizes to the mitochondria, but not Spn1-tdTomato in Tom20-GBP expressing cells. **F.** Spn3-GFP mislocalizes to the mitochondria, but not Rga4-mCherry, in Tom20-GBP expressing cells. **G.** Quantification of the coefficient of variation of Rga4-mCherry heterogeneity along cell sides in interphase cells, Spn3-GFP and Tom20-GBP expressing cells, as shown in E. [N=3, *n*=20-26 cells each, \**P*≤0.05, student’s t-test, scale bar=5μm].

If the septin cytoskeleton physically interacts with Rga4 and entraps it, this interaction should be detectable by measuring a change in Förster Resonance Energy Transfer (FRET) efficiency (Bajar et al., 2022; Bajar et al., 2016; McDonald et al., 2017). To test this, we utilized an Acceptor photobleach FRET method wherein FRET efficiency was measured between Rga4 and septin cytoskeletal proteins expressing GFP and RFP, respectively, prior to and following complete photobleaching of the red fluorescent acceptor (Liu et al., 2019). If interaction occurs between two proteins, bleaching of the red-tagged acceptor fluorophore should result in a nominal increase in GFP intensity, adjusted for normal photobleaching during the imaging process. If no interaction occurs, no increase in GFP intensity should be detectable following acceptor photobleaching. Indeed, when we look for interaction between two septin monomers in the cytoskeleton, Spn3-GFP and Spn1-tagRFP, we find a distinct increase in GFP intensity and FRET efficiency following acceptor photobleaching, indicating that there is interaction between these proteins (Supplementary Fig. S5A, B). We used this same approach to detect Spn1 and Rga4 interaction. We first identified regions of Rga4-GFP and Spn1-tagRFP overlap in the cell. Next, we bleached Spn1-tagRFP and monitored the intensity of Rga4-GFP. We fail to see an increase in FRET efficiency, and rather, Rga4-GFP intensity decreases to a degree consistent with regular photobleaching (Supplementary Fig. S5C, D). This absence of FRET efficiency increase between these two proteins suggests a lack of direct interaction between them.

Further, if the septin cytoskeleton physically interacts with and entraps Rga4, mislocalizing either the septin cytoskeleton or Rga4 should lead to mislocalization of the other. To test this, we used the GFP trap method with a GFP-binding protein (GBP) (Wang et al., 2016). We used the mitochondrial protein Tom20 tagged with GBP and RFP to entrap Spn3-GFP (Fig.6C). We find that in cells expressing Tom20-GBP-RFP, Spn3-GFP is no longer observed at the cortex, but rather appears along the mitochondria, colocalizing with Tom20. We asked if mislocalizing one of the components of the septin cytoskeleton could mislocalize the other components. We expressed Spn1-TdTomato and Spn3-GFP in cells expressing Tom20-GBP. We find that Spn3-GFP with Tom20-GBP at the mitochondria also entraps Spn1-tdTomato at that site (Fig.6D). This indicates that mislocalizing one of the septin components to the mitochondria is sufficient to mislocalize other proteins in the complex.

Next, we asked if Rga4-GFP can be mislocalized to the mitochondria with Tom20-GBP. We find that in cells expressing Tom20-GBP, Rga4-GFP appears away from the cortex and along the mitochondria (Fig.6E). However, in these cells, Spn1-tdTomato remained at the cortex and did not colocalize with the mitochondria and Rga4-GFP. Moreover, in Tom20-GBP expressing cells, Spn3-GFP at the mitochondria did not entrap Rga4-mCherry (Fig.6F). In these cells, Rga4-mCherry remained at the cortex along the cell sides. These observations, combined with our Acceptor Photobleach FRET data, suggest that the septin cytoskeleton does not regulate Rga4 distribution via physical interaction.

It is noteworthy that in the cells where the septin cytoskeleton localized to the mitochondria and away from the cortex, Rga4-mCherry at the cell sides appeared less punctate and more diffuse, similar to that observed in *spn1*Δ cells. To confirm this, we quantified the coefficient of variance of the Rga4-mCherry signal along the cell sides in control cells with those of cells co-expressing Tom20-GBP and Spn3-GFP. We find that the coefficient of variance of Rga4-mCherry is significantly reduced in cells where Spn3-GFP is mislocalized to the Tom20-GBP containing mitochondria (Fig.6F). This indicates that the absence of the septin cytoskeleton from the cell cortex is sufficient to alter Rga4 distribution pattern, even in cells where the septin proteins are expressed. These observations further support the hypothesis that the septin cytoskeleton at the cortex in interphase cells prevents Rga4 mobility along the cell sides.

### Septins promote the proper organization of membrane lipids

Our data show that the septin cytoskeleton does not physically interact with Rga4 to regulate its cortical dynamics and localization. Recent reports indicate that the septin cytoskeleton regulates membrane lipid organization in the budding yeast model system (El Alaoui et al., 2025; Pacheco et al., 2023). Similarly, septins in *Drosophila* modulate membrane PI(4,5)P_2_ resynthesis via regulation of the PIP5 kinase (Kumari et al., 2022). The septin cytoskeleton contains membrane-interacting domains, including an amphipathic helix that senses membrane curvature and specific interactions with membrane PI(4,5)P_2_ lipid (Beber et al., 2019; Bertin et al., 2010; Cannon et al., 2019; Lobato-Marquez et al., 2021; Woods et al., 2021; Zhang et al., 1999). Rga4 also contains membrane-interacting coiled-coil domains necessary for its membrane localization (Das et al., 2007). Thus, we asked if the septin cytoskeleton regulates Rga4 localization via membrane lipid organization. The ability of septin filaments to organize and exclude anionic and neutral lipids, respectively, has been shown to occur in budding yeast (El Alaoui et al., 2025). To study whether septins organize specific lipids along the *S. pombe* plasma membrane, we analyzed the localization of the fluorescently-tagged PI(4,5)P_2_ probe GFP-2x-PH_Plc_ in *spn1^+^* and *spn1*Δ cells (Willet et al., 2023). In *spn1^+^*cells, the PI(4,5)P_2_ biomarker, GFP-2x-PH_Plc_, is distributed primarily along the cell sides and non-growing cell ends, but is diminished at growing cell ends. However, in *spn1*Δ cells, GFP-2x-PH_Plc_ levels were increased all over the cortex, and particularly enhanced at the growing cell ends (Fig. 7A, B). It is possible that our observed increase in GFP-2x-PH_Plc_ at the cell ends of *spn1*Δ cells is the result of enhanced lipid binding capabilities due to the absence of the septin cytoskeleton, rather than an actual increase in the PI(4,5)P_2_ levels. If, on the other hand, the PI(4,5)P_2_ levels indeed increase, we would expect to see a decrease in the levels of its precursor PI(4)P. To test this, we analyzed the localization of the PI(4)P probe GFP-P4C-SidC in *spn1^+^* and *spn1*Δ cells. In *spn1^+^*cells, GFP-P4C-SidC localizes along the cell cortex, mainly concentrating at the growing cell ends (Snider et al., 2020). We find that, in *spn1*Δ cells, the overall PI(4)P levels at the cortex are reduced, including those at the cell ends (Fig. 7C, D). This suggests that the septin cytoskeleton regulates the PI(4,5)P_2_ levels and its organization at the plasma membrane. We asked if the disruption of the PI(4,5)P_2_ organization is due to changes in the kinase Its3 that converts PI(4)P to the PI(4,5)P_2_ (Deng et al., 2005). We find that, in *spn1^+^*cells, Its3-mNG is mainly localized along the cell sides and mostly excluded from the growing cell ends (Fig. 7E). In *spn1*Δ mutants, Its3-mNG levels at the cell ends are enhanced, in agreement with our data with the PI(4,5)P_2_ probe (Fig. 7E, F). Together, these data suggest that, in the absence of the septin cytoskeleton, the membrane lipid organization is disrupted, with excessive PI(4,5)P_2_ levels.

**Figure 7.**
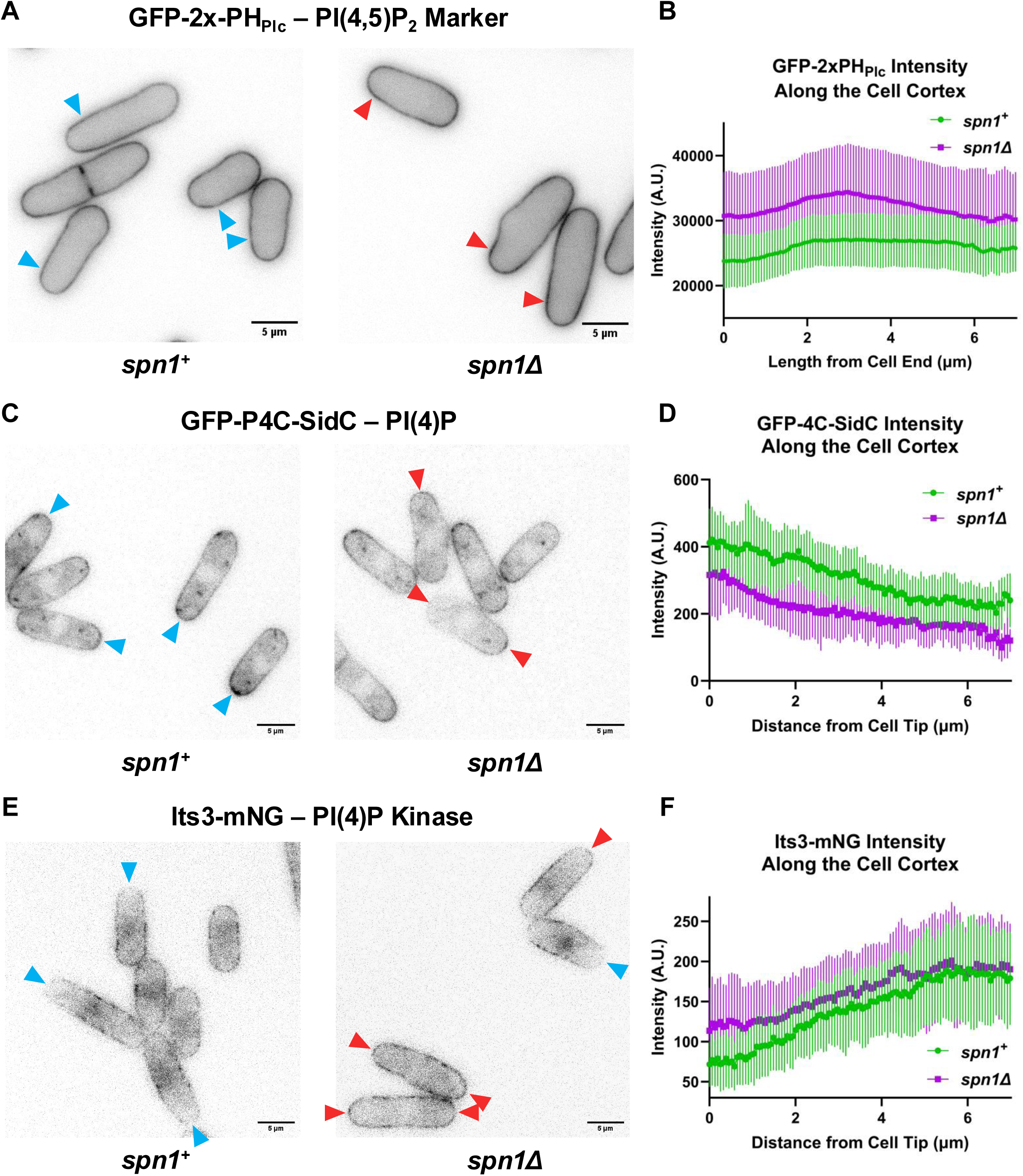
The septin cytoskeleton organizes lipids and lipid-associated enzymes along the cell membrane during interphase. **A.** Representative middle plane images of the PI(4,5)P_2_ fluorescent biomarker GFP-2x-PH_Plc_ in *spn1^+^* and *spn1*Δ cells. Blue and red arrowheads highlight decreased and increased GFP-2x-PH_Plc_ intensity at the border between the cell side and cell end, respectively. **B.** Quantification of GFP-2x-PH_Plc_ intensity along the cell cortex, measured from the very end of the cell to the middle. [N=3, *n*= 12-22 cells each, \*\*\*\**P*≤0.0001, student’s t-test]. **C.** Representative middle plane images of the PI(4)P fluorescent biomarker GFP-P4C-SidC in *spn1^+^* and *spn1*Δ cells. Blue and red arrowheads highlight increased and decreased GFP-P4C-SidC intensity at the cell ends, respectively. **D.** Quantification of GFP-P4C-SidC intensity along the cell cortex, measured from the very end of the cell to the middle. [N=3, *n*=12-20 cells each, \*\*\*\**P*≤0.0001, student’s t-test]. **E.** Representative middle plane images of the PI(4)P kinase Its3-mNG in *spn1^+^* and *spn1*Δ cells. Blue and red arrowheads highlight decreased and increased Its3-mNG intensity at the cell ends, respectively. **F.** Quantification of Its3-mNG intensity along the cell cortex, measured from the very end of the cell to the middle. [N=3, *n*=20-23 cells each, \*\*\*\**P*≤0.0001, student’s t-test, scale bar=5μm].

### Decreased PI(4,5)P_2_ levels at the cell membrane enhance Rga4-GFP membrane binding

Our data show that the loss of the septin cytoskeleton via *spn1*Δ results in increased PI(4,5)P_2_ levels at the cortex (Fig. 7A, B), while simultaneously leading to decreased distinct Rga4-GFP puncta formation and increased Rga4-GFP cortical spread, mobility, and cytoplasmic levels (Fig. 1D, E, G-J). We asked if a subsequent decrease in PI(4,5)P_2_ at the cell cortex would result in an opposite effect with regard to Rga4-GFP localization. To test this, we analyzed Rga4 localization in an *its3-1* temperature-sensitive mutant. When grown at the restrictive temperature of 36°C, *its3-1* expressing cells fail to convert PI(4)P to PI(4,5)P_2_ (Deng et al., 2005). At this high temperature, even in control cells, Rga4-GFP shows aberrant localization [data not shown], making characterization of Rga4 localization under these conditions difficult. We therefore examined Rga4 localization to the plasma membrane in the *its3-1* mutant at the semi-permissible temperature of 32°C over a 2-hour incubation time.

When observing Rga4-GFP localization in *its3-1* mutants grown under semi-restrictive conditions (32°C) for 2 hours, we find that Rga4-GFP more readily forms distinct puncta at the cell membrane (Fig. 8A, B). Furthermore, we see that with this increase in membrane puncta formation in the *its3-1* mutants, we subsequently observe an increase in the membrane-to-cytoplasmic ratio of Rga4-GFP in these cells, with a shift in the ratio favoring plasma membrane binding (Fig. 8 A-Middle Plane, C & D). These findings indicate that increased PI(4,5)P_2_ at the cell membrane resulting from disruption of the septin cytoskeleton in turn leads to increased Rga4-GFP mobility along the membrane and increased cytoplasmic exchange of Rga4. However, when the amount of PI(4,5)P_2_ at the cortex is reduced, Rga4-GFP more readily binds and forms distinct puncta on the membrane, with a subsequent decrease in the amount of Rga4 localized to the cytoplasm. Taken together, these results indicate that the septin cytoskeleton regulates PI(4,5)P_2_ organization at the plasma membrane, which in turn regulates Rga4 localization and polarity.

**Figure 8.**
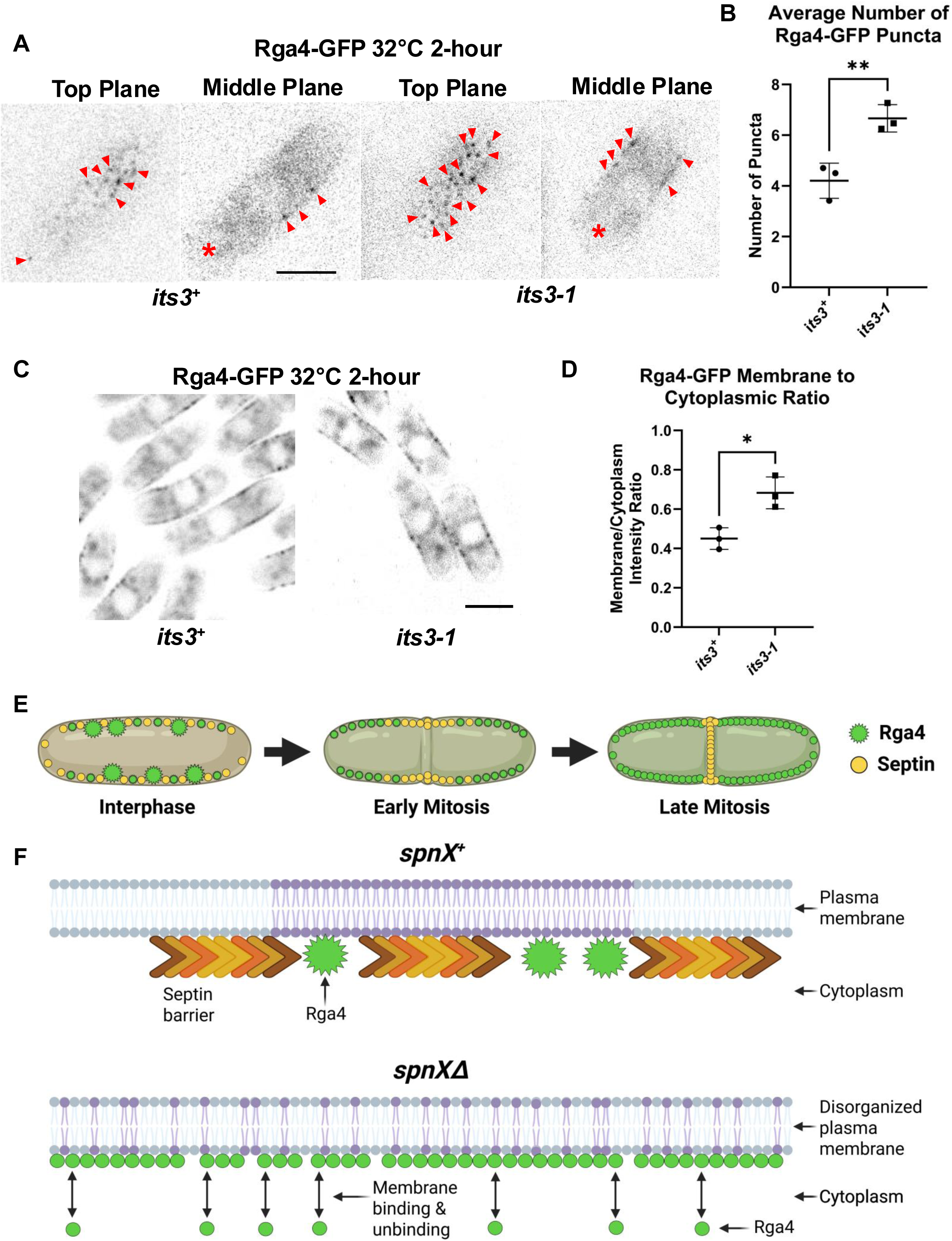
Disrupting PI(4,5)P_2_ levels at the cell cortex results in increased Rga4-GFP membrane binding and puncta formation. **A.** Representative images of top and middle plane images of Rga4-GFP puncta in *its3^+^* and *its3-1* cells grown at the semi-restrictive temperature of 32°C for 2 hours. Red arrows highlight Rga4-GFP puncta at the cell cortex. Red asterisks highlight differences in cytoplasmic Rga4-GFP intensity, particularly towards the cell ends. **B.** Quantification of the average number per cell of Rga4-GFP puncta at the cell cortex present in middle plane images of *its3^+^*and *its3-1* cells. [N=3, *n*>20, \*\**P*≤0.01, student’s t-test]. **C.** Representative images of Rga4-GFP in both *its3^+^* or *its3-1* cells growing at the semi-restrictive temperature of 32°C for 2 hours. Images consist of superimposed middle-plane images taken at 10 second intervals over a 5-minute time-lapse. **D.** Quantification of the membrane-to-cytoplasmic ratio of Rga4-GFP in *its3^+^* or *its3-1* cells grown for 2 hours at the semi-restrictive temperature of 32°C. [N=3, *n*>25, \**P*≤0.05, student’s t-test, scale bar=5μm]. **E.** Schematic model of Rga4 and septins in *S. pombe* throughout the cell cycle. Green dots indicate Rga4-GFP, and yellow dots indicate septins. Increased green highlighting of the cytoplasm denotes the increasing Rga4-GFP signal in the cytoplasm of *spn1^+^* cells throughout the cell cycle (Created with BioRender). **F.** Schematic model of the effects of the septin cytoskeleton on membrane lipid organization and Rga4 membrane localization. **Top.** Septin filaments help organize lipids along the plasma membrane, in turn facilitating immobile Rga4 puncta on the membrane. **Bottom.** The loss of the septin cytoskeleton results in disorganized membrane lipids, in turn resulting in a decreased ability of Rga4 to bind the membrane, instead promoting increased diffusion along the membrane, and increased exchange with the cytoplasm (Created with BioRender).

## Discussion

Our previous report showed that Cdc42 activation at the cell ends is cell-cycle-dependent and is turned off during cell division (Rich-Robinson et al., 2021). Cdc42 reactivation at the cell ends depends on the activation of the MOR/Orb6 kinase pathway and absence of Rga4 from the cell ends (Gupta et al., 2014; Rich-Robinson et al., 2021). Here we show that cell-cycle-dependent Rga4 localization is dependent on the septin cytoskeleton. The septin cytoskeleton at the cell sides prevents lateral diffusion of Rga4 along the membrane and keeps it away from the cell ends, where Cdc42 is activated for polarized growth. Once the cell enters mitosis, the septin cytoskeleton gradually drops from the cell sides to eventually form a medial ring, and this corresponds with Rga4 diffused localization to the cell ends, where Cdc42 is gradually inactivated (Fig. 8E).

After division in fission yeast, cells grow in a monopolar manner from their previously established old end and transition to bipolar growth when the <u>n</u>ew <u>e</u>nd takes <u>o</u>ff (NETO) (Mitchison and Nurse, 1985). Bipolar growth occurs when the old and new ends can compete with each other for Cdc42 activation (Das et al., 2012). The zone of growth at the cell ends is determined by the Cdc42 activation dynamics and the GAP Rga4 (Das et al., 2012; Das et al., 2007). Rga4 localizes to the cell sides like a corset and prevents the overflow of Cdc42 activation from the ends to these regions. This helps to maintain the cell width and proper polarity. During cell division, Rga4 localizes to the ingressing septum and prevents excessive Cdc42 activation at that site (Campbell et al., 2022). While Rga4 is mostly absent at the growing cell ends, in pre-NETO cells, Rga4 localizes to the non-growing ends. Recent reports show that Rga4 does not localize to the growing cell ends due to plasma membrane flows as a result of growth (Gerganova et al., 2021; Rutkowski et al., 2024). We find that in cells lacking the septin cytoskeleton, Rga4 levels at both cell ends increase due to increased mobility along the membrane and increased cytoplasmic exchange. However, the levels at the non-growing ends are higher than those at the growing ends, likely due to the effect of growth-dependent membrane flows. In these mutants, the cell ends receive increased levels of Rga4 due to diffusion, while the growth-dependent membrane flow removes Rga4.

Cdc42 activation dynamics at the cell ends depend on positive and time-delayed negative feedbacks (Das et al., 2012; Harrell et al., 2024; Howell et al., 2012). These feedbacks ensure that the two cell ends can sequentially activate Cdc42 to promote bipolar growth. Previous reports from our lab have shown that the GEF Gef1 is required for the establishment of Cdc42 activation at the new end and thus promotes bipolar growth (Hercyk and Das, 2019; Hercyk et al., 2019). Any disruption in the Cdc42 activation pattern results in its accumulation at one end at the expense of the other (Das et al., 2012; Harrell et al., 2024). We also observe this in the septin mutants, where Cdc42 dynamics are disrupted, resulting in monopolar growth. In *spn1^+^* cells, Rga4 can accumulate at the non-growing ends to some degree due to the lack of displacement from growth-dependent membrane flow. However, these levels are below a threshold and thus do not prevent subsequent Cdc42 activation at that site, allowing for bipolar growth. In septin mutants, however, the overall increased Rga4 accumulation at the cell ends, particularly at the non-growing end of monopolar cells, likely reaches a threshold. Once this threshold has been surpassed, relatively transient and low levels of Cdc42 activity at this end are not sufficient to effectively compete with the growing end for proper oscillatory dynamics and bipolar growth.

Rga4 contains coiled-coil domains that are required for its membrane localization (Das et al., 2007). In interphase cells, Rga4 mostly appears to form immobile puncta along the cell sides. During mitosis, these puncta disappear, and Rga4 displays faster dynamics with increased levels in the cytoplasm and diffuse localization along the cortex, including the cell ends. Our data indicate that the septin cytoskeleton maintains Rga4 puncta in interphase and restricts its localization to the cell sides. However, this regulation is not mediated via protein-protein interaction. Due to the geometry of septin filament organization during interphase along the long axis of the cell, it is also unlikely that the septin cytoskeleton restricts Rga4 diffusion along the plasma membrane via physical obstruction. Our data show that, in the fission yeast system, the septin cytoskeleton plays a role in affecting plasma membrane lipid organization during interphase. The presence of septins appears to promote the proper localization and amounts of PI(4)P and PI(4,5)P_2_ along the plasma membrane of interphase cells. This localization coincides with, and is likely downstream of, the localization of the PI(4)P kinase Its3, which is also reliant on the septin cytoskeleton for its proper localization during interphase. In keeping with this observation, we find that in cells with diminished PI(4,5)P_2_ levels, Rga4 membrane puncta and levels are enhanced. Thus, we posit that the septin cytoskeleton limits membrane PI(4,5)P_2_ levels to promote Rga4 localization to immobile puncta along the cell sides. In the absence of the septin cytoskeleton, membrane PI(4,5)P_2_ levels are enhanced, and Rga4 dynamics and exchange with the cytoplasm increase (Fig. 8F).

The septin cytoskeleton in fission yeast has been investigated mostly in the context of cytokinesis (An et al., 2004; Berlin et al., 2003; Martin-Cuadrado et al., 2005; Munoz et al., 2014; Singh et al., 2024; Wang et al., 2015). In budding yeast, the septins organize as a collar around the bud neck and form a ring during cytokinesis (Bridges and Gladfelter, 2015; Woods and Gladfelter, 2021). The septin collar in budding yeast has been shown to restrict GAP localization near the bud neck to promote bud emergence for daughter cell formation (Okada et al., 2013). This indicates that restricting GAP localization by septin filaments is a conserved phenomenon. Here, we show that the septin cytoskeleton forms short linear filaments along the long axis of the cell during interphase. While the core septins Spn1 and Spn4 are critical for the medial ring formation during cytokinesis, all 4 septin proteins Spn1-4 are required for their role in cell polarity. This indicates that the nature of the filaments formed in interphase is distinct from that during mitosis. In fission yeast, the septin ring forms as anaphase B ends, when the actomyosin ring starts to constrict. However, the septin filaments at the cell sides begin to disappear at the onset of mitosis, well before the septin ring forms. The septin proteins have membrane-interacting domains and are involved in lipid metabolism (Mela and Momany, 2022; Spiliotis and McMurray, 2020; Woods and Gladfelter, 2021). A mitotic signal may disable the septin-membrane interaction, resulting in its loss from the cell sides. In budding yeast, the LKB1-like kinase Elm1 has been shown to promote septin hourglass assembly and stability during bud emergence (Marquardt et al., 2020). In mitosis, Elm1 regulates the Nim1/PAR-1-related kinase Gin4 kinase to enable septin double ring formation (Marquardt et al., 2024). However, the Gin4 homolog in fission yeast, Cdr2, does not regulate septin ring formation (Morrell et al., 2004). Further investigation will determine what leads to the loss of septins from the cell sides in mitosis and its subsequent ring formation after anaphase B.

Our findings here show a role for the septin cytoskeleton in regulating polarized cell growth in a cell-cycle-dependent manner. While most cell-cycle-dependent regulation is mediated via kinase-substrate phosphorylation (Kaldis, 1999; Malumbres, 2014; McCusker and Kellogg, 2012), here we show a novel regulatory system where Rga4 localization and dynamics are indirectly controlled by regulating the distribution of specific lipids along the plasma membrane via septin localization during interphase. While the septin rings have been mostly identified in fungal cells, it is possible that septins in other eukaryotes also undergo cell-cycle-dependent changes in their organization (Delic et al., 2024; Marquardt et al., 2021). In several eukaryotes, septins have been shown to act as diffusion barriers and maintain asymmetry in the cell (Caudron and Barral, 2009; Spiliotis and McMurray, 2020). Cell-cycle-dependent changes in septin organization may alter their ability to act as barriers to the movements of both proteins and membrane lipids.

## Methods and Materials

### Strains and cell culture

The *S. pombe* strains used in this study are listed in Table 1. All strains are isogenic to the original strain PN567. Cells were cultured in yeast extract with supplements (YES – Sunrise Science Products) medium and grown exponentially at 25°C, unless specified otherwise. Standard techniques were used for genetic manipulation and analysis (Moreno et al., 1991). Cells were grown exponentially for at least three rounds of eight generations each before imaging.

### Microscopy

Imaging was performed at room temperature (23–25°C). We used a spinning disk confocal microscope system with a Nikon Eclipse inverted microscope with a 100×/1.49 NA lens, a CSU-22 spinning disk system (Yokogawa Electric Corporation), and a Photometrics EM-CCD camera (Photometrics Technology Evolve with excelon Serial No: A13B107000). Images were acquired with MetaMorph (Molecular Devices, Sunnyvale, CA).

A Nikon Ti2 Eclipse wide-field microscope was also used with a 100x/1.49 NA objective, an ORCA-FusionBT digital camera (Hamamatsu Model: C15440-20UP Serial No: 500428, Hamamatsu, Hamamatsu City, Japan). Images were acquired using Nikon NIS Elements (Nikon, Melville, NY). Fluorophores were excited using an AURA Light Engine system (Lumencor, Beaverton, OR).

Additionally, microscopy was performed with a 3i spinning disk confocal using a Zeiss AxioObserver microscope with integrated Yokogawa spinning disk (Yokogawa CSU-X1 A1 spinning disk scanner) and a 100x/1.49 NA objective. Images were acquired with a Teledyne Photometrics Prime 95b back-illuminated sCMOS camera (Serial No: A20D203014, Tucson, AZ). Images were acquired using SlideBook (3i Intelligent Imaging innovations, Denver, CO).

### Fluorescence Intensity Acquisition & Quantification

Image analysis was performed on a Windows computer using the public domain NIH ImageJ program (developed at the U.S. National Institutes of Health and available on the Internet at http://rsb.info.nih.gov/nih-image/). Images were acquired at either single, mid-section focal planes, or as Z-stacks with 0.2-0.4 μm focal plane intervals. Z-stack images were superimposed using SUM-projection for analysis. Mean intensity measurements were used for analysis, along with integrated density values where specified. Prior to intensity acquisition, a region of interest devoid of cells was used for background subtraction.

### Analysis of protein localization heterogeneity along the cell sides

Heterogeneity of protein localization along the cell sides was analyzed by measuring the coefficient of variance of protein distribution along said sides. Line profiles of each cell side were drawn using ImageJ, and the standard deviations and averages of fluorescence intensity along these line profiles were measured. The coefficient of variance was obtained by dividing the measured standard deviation by the obtained average, and then multiplying the result by 100%.

### Fluorescence Recovery After Photobleaching

Fluorescence recovery after photobleaching (FRAP) was performed as described before (Onwubiko et al., 2019). Fluorescently tagged proteins along the cell cortex were first focused at the medial z-plane, and an ROI was selected for bleaching. The ROI was imaged 10 times pre-bleaching and then bleached for 10 s, and images were then acquired every 3 s for 50–100 images. A non-bleached spot on the cortex for the same cell was analyzed to account for loss of signal due to photobleaching over time. A cell-free region of the image was used for background subtraction. Intensity values for each ROI were corrected for loss of signal over time and background signal. The intensity values were then normalized against the mean pre-bleach value to determine the recovery fraction over time. At least ten cells were analyzed for each genotype for each experiment, and carried out in triplicate. The average corrected and normalized intensity values were then plotted as a fraction of recovery over time. The fraction of recovery over time was plotted and smoothed using GraphPad *Prism*. The half-life (t_1/2_) was then computed as the time for recovery of 50% of the signal.

### Acceptor-photobleach Förster Resonance Energy Transfer (FRET)

FRET imaging was performed using a 3i spinning disk confocal using a Zeiss AxioObserver microscope with integrated Yokogawa spinning disk (Yokogawa CSU-X1 A1 spinning disk scanner) and a 100x/1.49 NA objective. Images were acquired with a Teledyne Photometrics Prime 95b back-illuminated sCMOS camera (Serial No: A20D203014, Tucson, AZ). Images were acquired using SlideBook (3i Intelligent Imaging innovations, Denver, CO). An acceptor photobleaching method was employed for FRET imaging in live cells. The Spn1-tagRFP fluorophore was bleached throughout the cells using 40% 561 nm laser power for 3 cycles. Middle plane images in the donor GFP channel were acquired pre- and post-bleach. FRET percentages were calculated by first correcting for background and photobleaching between acquisitions pre- and post-bleach, and subsequently calculating the percentage change in GFP donor fluorescence at areas of donor and acceptor fluorophore colocalization along the cell cortex using ROIs of a consistent size.

## Supplemental information titles and legends

Document S1. Figures S1–S5 and Table S1

## Data Availability

The data generated are available in the published article and its online supplemental material.

## Acknowledgements

We thank Kathy Gould, Fulvia Verde, Pilar Perez, Chuanhai Fu, Jian-Qiu Wu, and Damian Brunner for providing strains and Bret Judson at the Boston College Imaging Core for imaging support. This work is supported by the National Institutes of Health grant R01GM136847 to M.D.

## Author contributions

Conceptualization: M.D. and J.M.; Methodology: M.D. and J.M.; Validation: M.D. and J.M.; Formal analysis: M.D. and J.M.; Investigation: M.D., J.M., G.C.; Resources: M.D.; Data curation: J.M. and G.C.; Writing-original draft: M.D. and J.M.; Writing - review & editing: M.D.; Supervision: M.D.; Project administration: M.D.; Funding acquisition: M.D.

