## Supplementary Material for "Septins mediate cell-cycle-dependent exclusion of Cdc42 GAP Rga4 from growth sites in fission yeast"

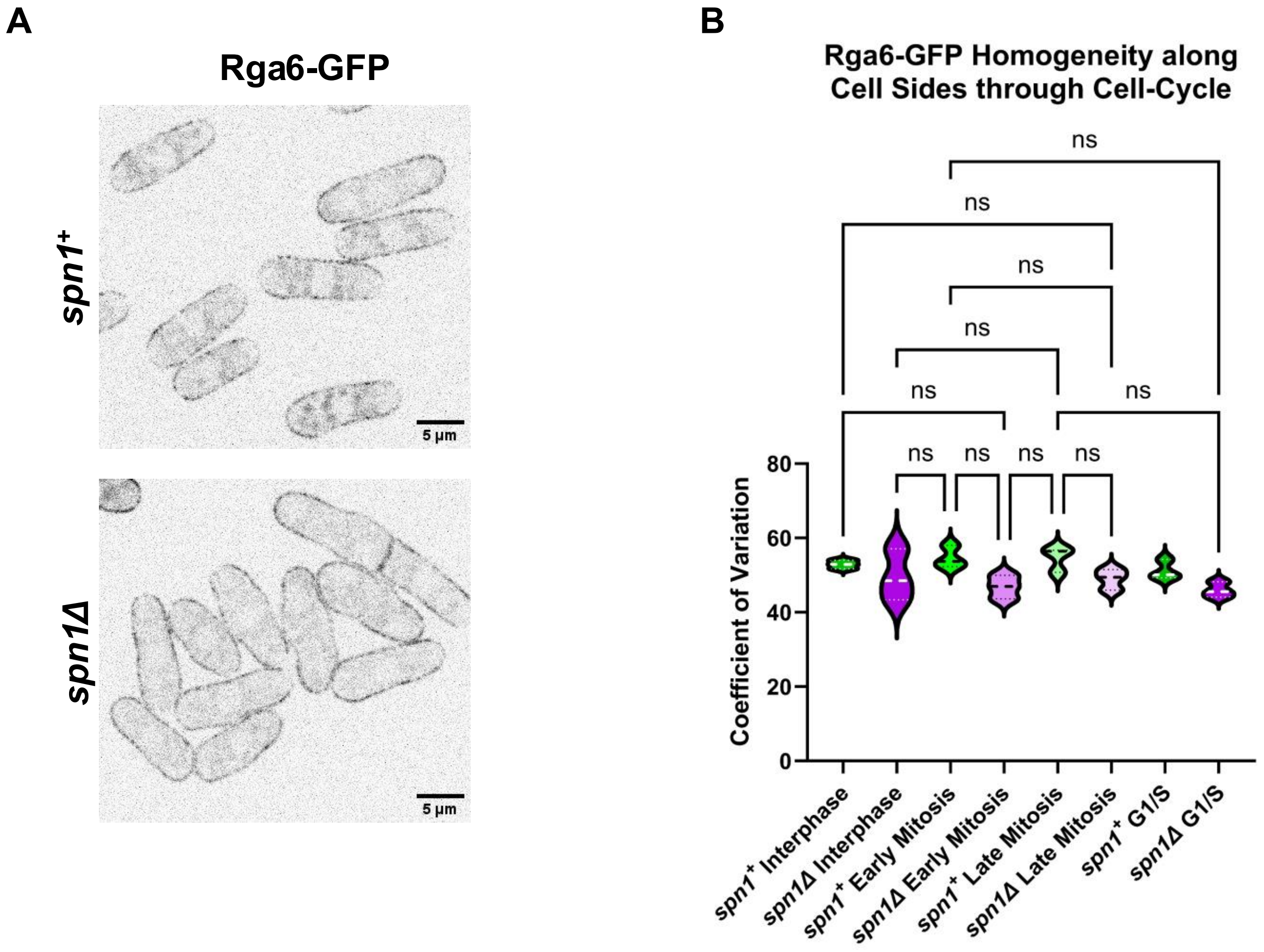


**Supplementary Figure 1. The Septin cytoskeleton does not disrupt Rga6 localization pattern along the cell sides. A.** Rga6-GFP localization in *spn1*$^{+}$ and *spn1Δ* cells. **B.** Quantification of the coefficient of variation of Rga6-GFP distribution through the cell-cycle in *spn1*$^{+}$ and *spn1Δ* cells. [N=3, n=40-50 cells each, ns=not significant, one-way ANOVA, scale bar=5μm].

### Supplementary Figure 2.

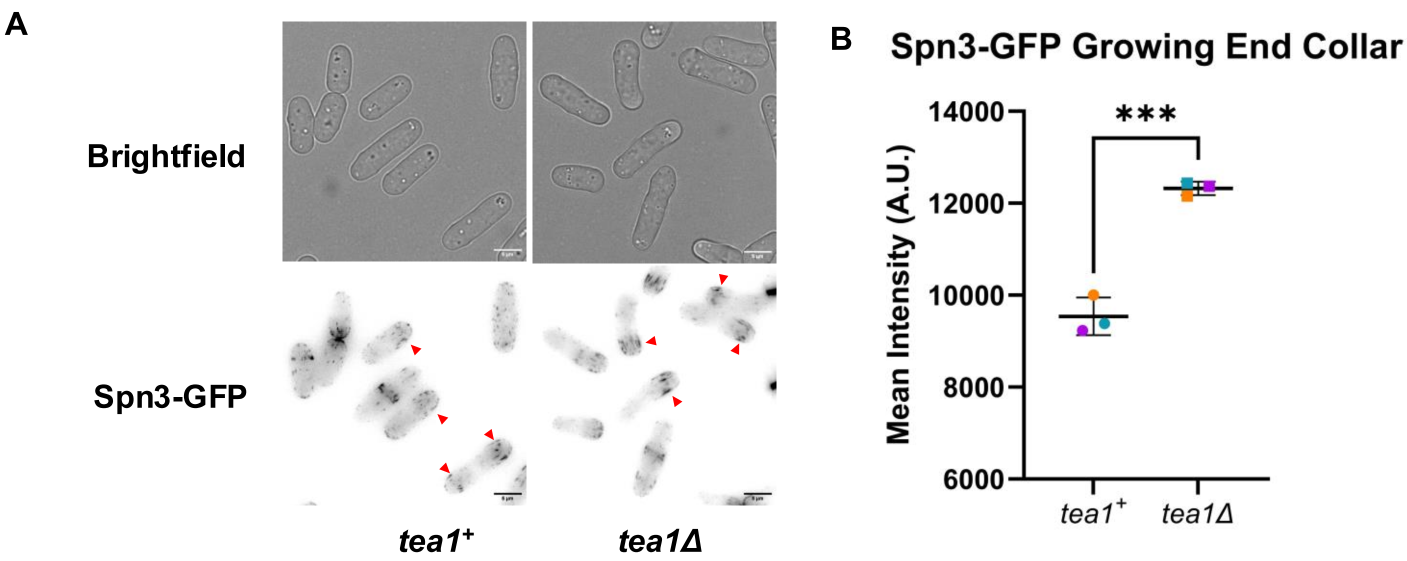


**Supplementary Figure 2. Septin filaments are enriched near the growing cell ends. A.** Brightfield images of *tea1*$^+$ and *tea1Δ* cells displayed above. Fluorescent images of Spn3-GFP displayed below. **B.** Quantification of Spn3-GFP intensity at growing cell end collars, near the cell end. [N=3, *n*=15-20 cells each, ****P* ≤0.001, student's t-test]. Max Projection images used for representation. Sum Projection images used for quantification. Scale bar=5 μm.

**Supplementary Figure 3.**

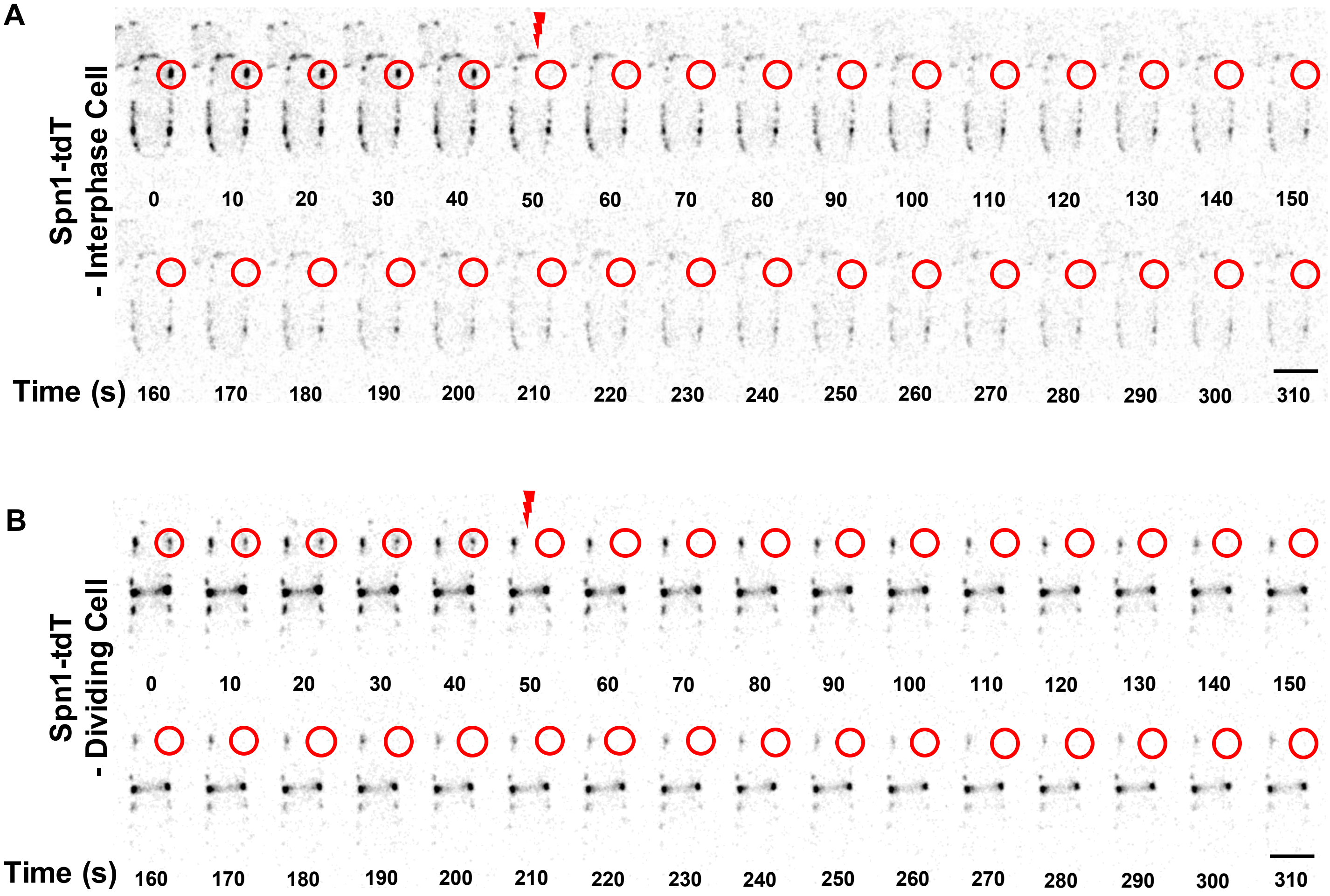


**Supplementary Figure 3. Septin filaments are stable at the cell cortex. A-B.** Timelapse of Spn1-tdTomato expressed in interphase **(A)** and dividing cells **(B)** after photobleaching. Red lightning marks the timing of bleaching and red circle marks zone of bleach. Scale bar, 5μm

### Supplementary Figure 4.

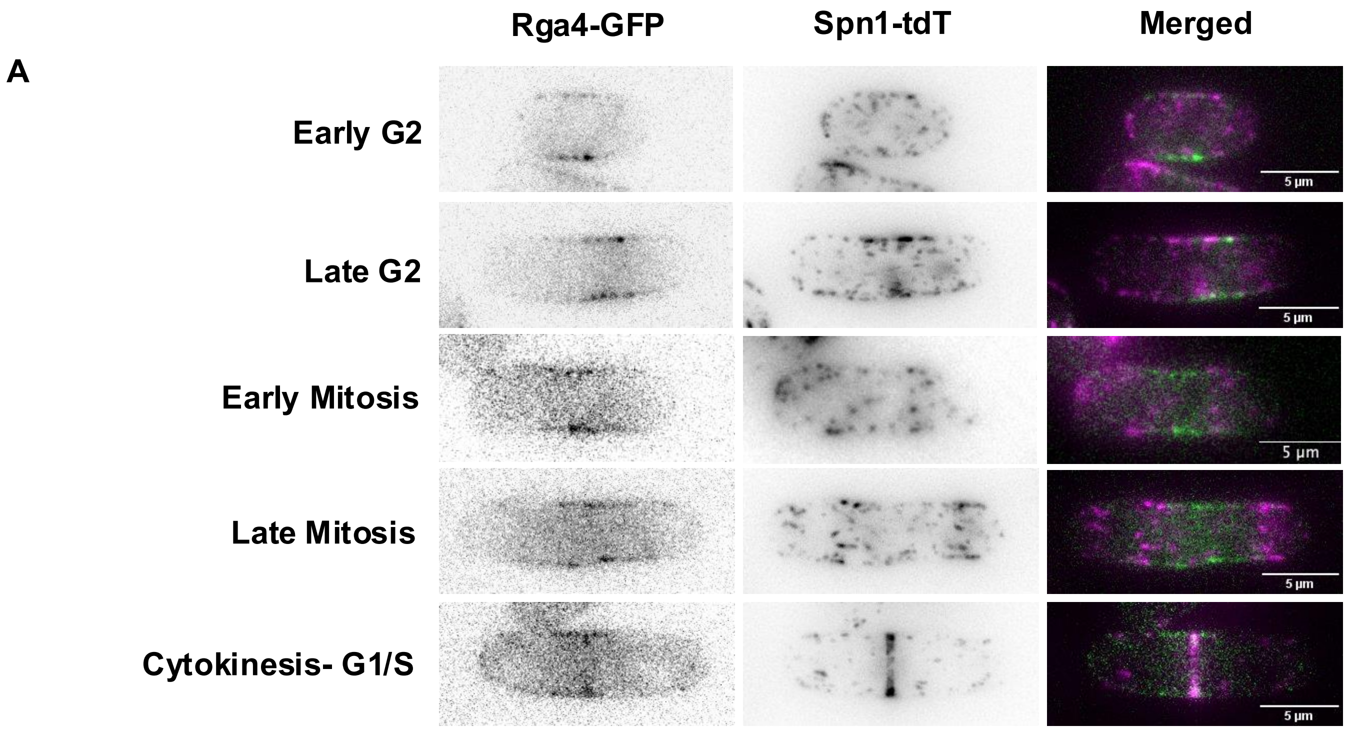


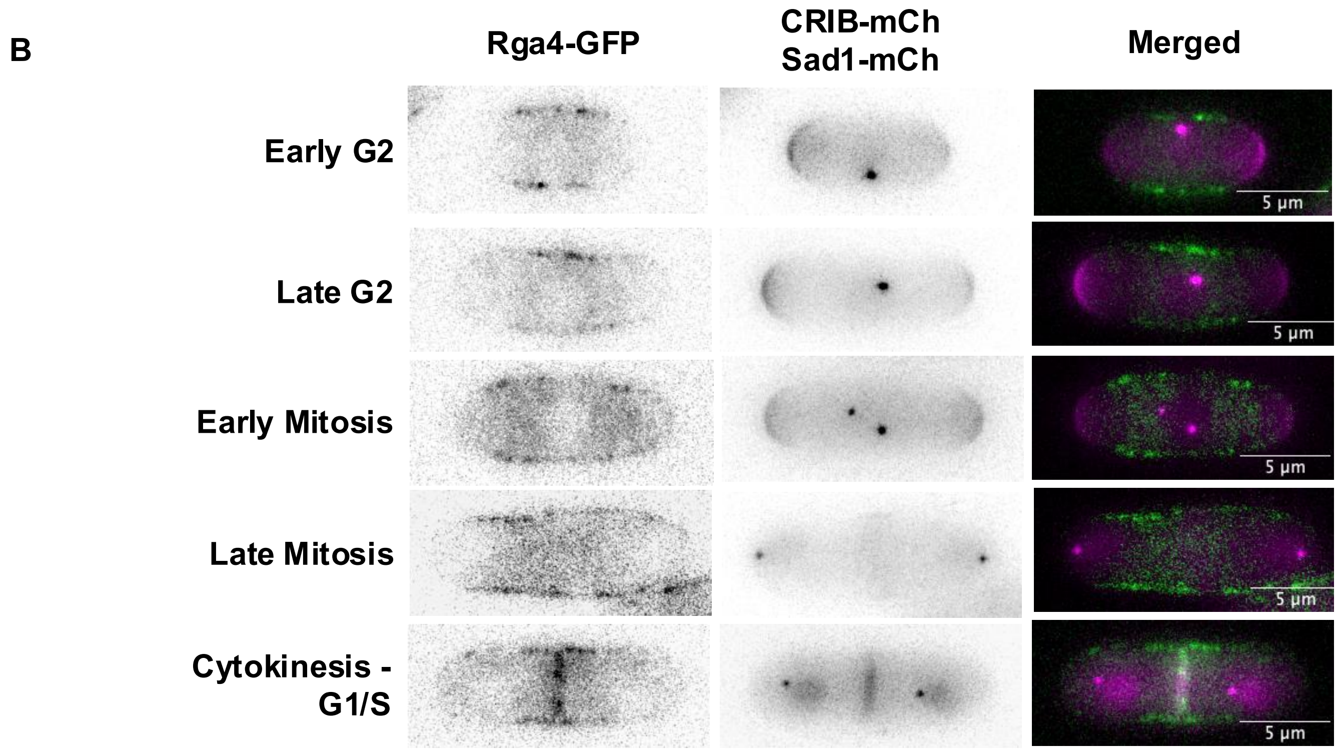


**Supplementary Figure 4. As septins recede from the cell sides, Rga4 spreads out along the membrane and Cdc42 activity diminishes at the cell ends. A.** Representative middle-plane images of Rga4-GFP and Maximum Z-Projections of Spn1-tdTomato images throughout the cell-cycle. **B.** Representative middle-plane images of Rga4-GFP and Maximum Z-Projections of the active Cdc42 biomarker CRIB-mCherry and spindle pole body marker Sad1-mCherry throughout the cell cycle. Scale bar=5µm

### Supplementary Figure 5.

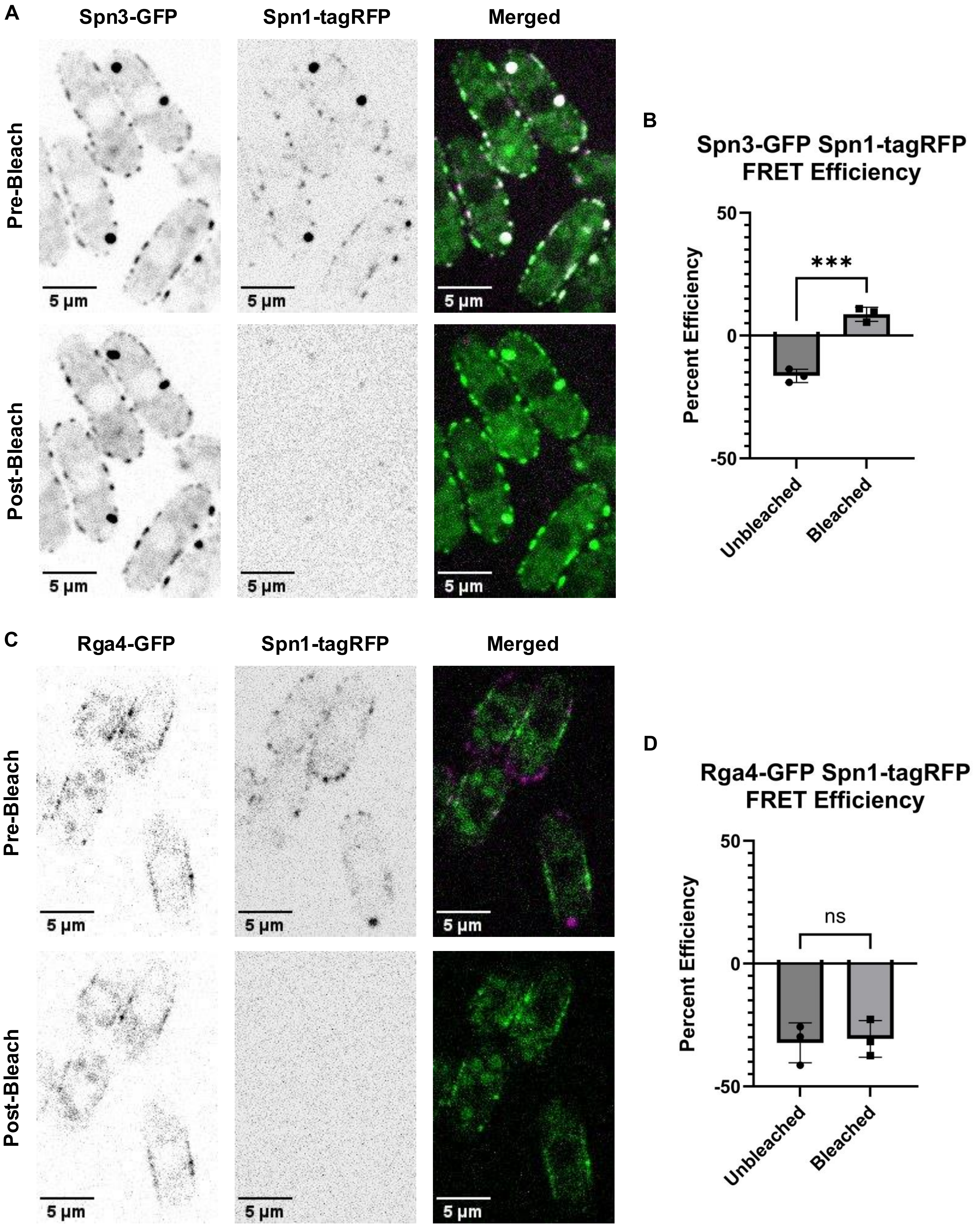

**Supplementary Figure 5. Acceptor bleach FRET analysis indicates that Rga4 does not interact with the septin cytoskeleton (Spn1-tagRFP specifically). A.** Representative fluorescent images of Spn3-GFP and Spn1-tagRFP expressing cells before and after photobleaching Spn1-tagRFP. **B.** Quantification of the percent FRET efficiency before and after Spn1-tagRFP photobleaching. **C.** Representative fluorescent images of Rga4-GFP and Spn1-tagRFP expressing cells before and after photobleaching. **D.** Quantification of the percent FRET efficiency before and after Spn1-tagRFP photobleaching. [N=3, *n*=16-24 cells each , ****P* ≤0.001, ns=not significant, student's t-test, scale bar=5μm].

| Strain name | Mating | Genotype | Source |
| --- | --- | --- | --- |
| FV781 | h+ | *rga4-GFP::kanMX ura4-D18 ade6*$^{-}$ *leu1-32* | Das et al., 2007 |
| FV513 | h- | *rga4Δ::ura4+ade6-704leu1-32ura4-D18* | Das et al., 2007 |
| FV467 | h+ | *tea1Δ::ura4*$^{+}$ *ura4-D18 ade6-M210* | Verde et al., 1995 |
| PN567 | h+ | *leu1-32 ade6-m210 ura4-D18* | Paul Nurse |
| PPG56.68 | *h+* | *scd2-GFP::kanMX leu1-32 ura4-d18* | Rincon et al., 2007 |
| CF 4473 | h- | *spn1-tdTomato::Nat*$^{R}$ *: leu1-32 ade6-m210 ura4-D18* | Zheng et al., 2022 |
| CF 4644 | h+ | *spn4-tdTomato::Nat*$^{R}$ *myo2-GFP:ura4*$^{+}$ *sid4-GFP:kanMx ade6-210 leu1-32 ura4-D18* | Zheng et al., 2023 |
| DB 4956 | h- | *spn1-tagRFP::kanR spn3-GFP::kanR* | Liu et al., 2019 |
| JW709 | h- | *spn3-Δ2::kanMX6 ade6-M210 leu1-32 ura4-D18* | Singh et al., 2025 |
| JW1091 | h- | *spn1-mEGFP-kanMX6 ade6-M210 leu1-32 ura4-D18* | Gregory et al,. 2025 |
| JW | h- | *spn4-tdTomato-natMX6 ade6-M210 leu1-32 ura4-D18* | Gregory et al,. 2025 |
| JW9724 | h- | *spn4-Δ2::hphMX6 ade6-210 ura4-D18 leu1-32* | Singh et al., 2025 |
| JW9732 | h- | *spn2-Δ1::hphMX6 ade6-210 ura4-D18 leu1-32* | Singh et al., 2025 |
| KGY243-2 | h- | *its3-mNG:hygR ade6-M210 leu1-32 ura4-D18* | Snider et al., 2020 |
| KGY269-2 | h- | *GFP-2xPHPlc:leu*$^{+}$ *ade6-M210 leu1-32 ura4-D18* | Snider et al., 2020 |
| KGY3244 | h- | *spn3-GFP::kanMx leu1-32 ura4-D18 ade6-m210* | An et al., 2004 |
| KGY4230 | h- | *spn2-GFP::kanMx leu1-32 ura4-D18 ade6-m210* | An et al., 2005 |
| KGY19217 | h+ | *GFP-P4CSidC:leu*$^{+}$ *ade6-M210 leu1-32 ura4-D18* | Snider et al., 2020 |
| YSM1649 | h- | *its3-1 leu1-32 ura4-D18* | Zhang et al., 2000 |
| YMD754 | h+ | *rga4Δ::ura4*$^{+}$ *CRIB-3xGFP:ura4*$^{+}$ *rlc1-tdTomato::NatR sad1-mCherry::kanMX ade6-704 leu1-32 ura4-D18* | This work |
| YMD771 | h- | *rga6-GFP::kanMX rlc1-tomato::Nat*$^{R}$ *sad1-mCherry::kanMX leu1-32 ura4-d18* | This work |
| YMD772 | | *rga4-GFP::kanMX rlc1-tdTomato::Nat*$^{R}$ *sad1-mCherry::KanMx* | This work |
| YMD1192 | h- | *scd1-mNeonGreen::kanMX ura4-D18 ade6-leu1-32* | This work |
| YMD1551□ | | *CRIB-3xGFP:ura4*$^{+}$ *rlc1-tdTomato::Nat*$^{R}$ *sad1-mCherry::kanMX* | This work |
| YMD1684 | h- | *spn1Δ::ura4*$^{+}$ *ura4-D18 leu1-32* | Rich-Robinson et al., 2021 |
| YMD1686 | h+ | *spn1Δ::ura4*$^{+}$ *ura4-D18 leu1-32* | Rich-Robinson et al., 2021 |

| | | | |
|---|---|---|---|
| YMD1795 | | *rga4-GFP::kanMX spn1Δ::ura4$^{+}$ ura4-D18 leu1-32* | This work |
| YMD1853 | | *spn1Δ::ura4$^{+}$ rga4-GFP::KanMX rlc1-tdTom::Nat$^{R}$ Sad1-mCherry::kanMx* | This work |
| YMD1880 | | *spn3-GFP::kanMx CRIB-mCherry::leu1$^{+}$ sad1-mCherry::kanMX leu1-32 ura4-D18 ade6-m210* | This work |
| YMD1921 | | *spn1Δ CRIB-GFP Rlc1-tdTomato Sad1-mCherry #4* | This work |
| YMD2156 | | *spn1Δ::ura4$^{+}$ rga6-GFP::kanMX Rlc1-td tomato::Nat$^{R}$ Sad1-mCherry::kanMX* | This work |
| YMD2365 | h- | *rga4-GFP::kanMX spn1-tagRFP::kanR* | This work |
| YMD2422 | | *spn1Δ::ura4$^{+}$ spn4-tdTomato::Nat$^{R}$ ura4-D18 ade6-M210* | This work |
| YMD2423 | | *spn1Δ::ura4$^{+}$ spn3-GFP::kanMx ura4-D18 ade6-M210* | This work |
| YMD2424 | | *spn1Δ::ura4$^{+}$ spn2-GFP::kanMx ura4-D18 ade6-M210* | This work |
| YMD2450 | h+ | *rga4-GFP::kanMX spn1-tdTomato::Nat$^{R}$ ura4-D18 ade6- leu1-32* | This work |
| YMD2522 | h- | *spn3-GFP::kanMx tea1Δ::ura4$^{+}$ ura4-D18 ade6-M210* | This work |
| YMD2544 | | *spn3-GFP::kanMX spn1-tdTomato::Nat$^{R}$ ura4-D18 ade6- leu1-32* | This work |
| YMD2576 | | *spn3-GFP::kanMx tom20-GBP-RFP::Hph leu1-32 ura4-D18 ade6-m210* | This work |
| YMD2578 | | *rga4-GFP::kanMX spn1-tdTomato::Nat$^{R}$ tom20-GBP::Hph ura4-D18 ade6- leu1-32* | This work |
| YMD2579 | | *spn3-GFP::kanMx tom20-GBP-RFP::Hph rga4-mCherry::kanMX ura4-D18 ade6- leu1-32* | This work |
| YMD2583 | | *rga4-GFP::kanMX CRIB-mCherry::leu1$^{+}$ sad1-mCherry::kanMX ura4-D18 ade6- leu1-32* | This work |
| YMD2626 | | *spn1Δ::ura4$^{+}$ GFP-2xPHPlc:leu+ ade6-M210 leu1-32 ura4-D18* | This work |
| YMD2628 | | *spn1Δ::ura4$^{+}$ its3-mNG:hygR ade6-M210 leu1-32 ura4-D18* | This work |
| YMD2699 | | *spn3-GFP::kanMX spn1-tdTomato::Nat$^{R}$ Tom20-GBP::Hph ura4-D18 ade6- leu1-32* | This work |
| YMD2837 | | *spn1Δ::ura4$^{+}$ GFP-P4CSidC:leu+ ade6-M210 leu1-32 ura4-D18* | This work |
| YMD 2842 | | *rga4Δ::ura4$^{+}$ spn1Δ::ura4$^{+}$ CRIB-3xGFP:ura4$^{+}$ rlc1-tdTomato::NatR sad1-mCherry::kanMX ade6-704 leu1-32 ura4-D18* | This work |

| | | |
|---|---|---|
| YMD 2960 | *its3-1 rga4-GFP::kanMX leu1-32 ura4-D18* | This work |
